# Novel multiparametric bulk and single extracellular vesicle pipeline for adipose cell-specific biomarker discovery in paired human biospecimens

**DOI:** 10.1101/2024.04.18.590172

**Authors:** Mangesh Dattu Hade, Jacelyn Greenwald, Paola Loreto Palacio, Kim Truc Nguyen, Dharti Shantaram, Bradley L. Butsch, Yongseok Kim, Sabrena Noria, Stacy A. Brethauer, Bradley J. Needleman, Willa Hsueh, Vicki H. Wysocki, Eduardo Reátegui, Setty M. Magaña

## Abstract

Obesity remains a growing and global public health burden across a broad spectrum of metabolic, systemic, and neurodegenerative diseases. Previously considered merely a fat storage depot, adipose tissue is now recognized as an active endocrine organ crucial for metabolic and systemic regulation of local and distant organs. A burgeoning line of investigation centers on adipose-derived extracellular vesicles (ADEVs) and their pivotal role in obesity-associated pathobiology. However, robust methodologies are lacking for specifically isolating and characterizing human ADEVs. To bridge this gap, we have developed a robust multiparametric framework incorporating bulk and single EV characterization, proteomics, and mRNA phenotyping. EVs from matched human visceral adipose tissue, mature adipocyte-conditioned media, and plasma collected from the same individual bariatric surgical patients were analyzed and subjected to bottom-up proteomics analysis. This framework integrates bulk EV proteomics for cell-specific marker identification and subsequent single EV interrogation with single-particle interferometric reflectance imaging (SP-IRIS) and total internal reflection fluorescence (TIRF) microscopy. Our proteomics analysis revealed 76 unique proteins from adipose tissue-derived EVs (ATEVs), 512 unique proteins from adipocyte EVs (aEVs), and 1003 shared proteins. Prominent pathways enriched in ATEVs included lipid metabolism, extracellular matrix organization, and immune modulation, while aEVs exhibited enhanced roles in chromatin remodeling, oxidative stress responses, and metabolic regulation. Notably, adipose tissue-specific proteins such as adiponectin and perilipin were highly enriched in ADEVs and confirmed in circulating plasma EVs. Colocalization of key EV and adipocyte markers, including CD63 and PPARG, were validated in circulating plasma EVs. In summary, our study paves the way toward a tissue and cell-specific, multiparametric framework for an ‘adiposity EV signature’ in obesity-driven diseases.

## Introduction

Obesity has emerged as one of the most critical global health crises, reaching epidemic proportions and straining public health systems worldwide^1^. Obesity’s association with a myriad of diseases, including metabolic disorders, cancer, autoimmunity, and neurodegenerative diseases, underscores the urgency of understanding the underlying pathophysiology driving these conditions^2–6^. Once considered merely an inert energy storage tissue, adipose tissue is now recognized as an active endocrine organ that regulates metabolic homeostasis, immune responses, and systemic health^7^. The dysfunction of adipose tissue, marked by chronic inflammation, altered adipokine secretion, and impaired adipocyte differentiation, is a hallmark of obesity and contributes significantly to developing obesity-related comorbidities^8,9^.

Recent advances in extracellular vesicle (EV) research have shed light on the broad translational potential of EVs as pathogenic drivers of disease, diagnostic and prognostic biomarkers, and cell-free, therapeutic vectors in a myriad of diseases^10^. EVs exert phenotypic and genotypic changes in recipient cells by transmitting their cell-specific and context-specific bioactive cargoes (e.g., proteins, lipids, and nucleic acids)^11–13^. Circulating plasma EVs (bEVs) represent heterogenous EV subpopulations released by hematopoietic cells, as well as tissue-derived EVs. Adipose-derived EVs (ADEVs) are estimated to account for more than 80% of the tissue-derived, circulating EV plasma pool and account for the majority of EV-associated miRNA in obesity models^14^. These recent findings have positioned ADEVs as novel and central players in the pathogenesis of obesity-driven diseases.

Two key subsets of ADEVs include adipose tissue-derived EVs (ATEVs) and mature adipocyte-derived EVs (aEVs), each with unique pathobiological roles and cell-specific phenotypic repertoires. ATEVs, consisting of EVs originating from a heterogenous composite of cells from the vascular stroma, such as pre-adipocytes, immune cells, and support cells, reflect the multicellular adipose tissue microenvironment^11,15^. On the other hand, EVs released from mature adipocytes provide insights into the molecular messages adipocytes utilize to communicate with other cells and tissues^16^. Despite the growing interest in ADEVs, the full translational potential of human-sourced EVs is hindered by several methodological and biological challenges, including a lack of standardized, reproducible, and robust isolation and cell-specific characterization protocols^17,18^. While a handful of studies have investigated human ADEVs^12,19,20^, most ADEV studies have focused on rodent models^21–23^, despite recognized cross-species differences in adipose markers between rodents and humans^24^. This cross-species variation highlights the critical need for human-based, obesity EV paradigms.

To address the current technical and biological challenges impeding the clinical translation of human ADEVs, we developed a novel, comprehensive, human-based, multiparametric framework for isolating and characterizing ADEVs. Advanced, high-resolution technologies were employed to characterization EVs from human visceral adipose tissue, cultured mature adipocyte-conditioned media, and plasma derived from the same bariatric individual. while detailed proteomic profiling of adipose EVs was performed to facilitate the development of an adipose EV-based liquid biopsy. Critically, tissue-and cell-specific adipose EV signatures were validated in circulating plasma EVs from the same bariatric patients. Finally, in silico functional prediction analysis was conducted to provide insights into the potential pathobiological role ADEVs in metabolic disease.

## Results

### Multiparametric isolation and characterization of obesity-associated EVs

Most human ADEV biomarker studies have focused on biofluids or in vitro ADEV sources; however, technical challenges must be overcome for tissue-derived EVs to maintain EV integrity^25–27^. To ensure the integrity of tissue-derived adipose EVs, we employed comprehensive bulk and single-vesicle characterization methods (Figure 1).

**Figure 1.**
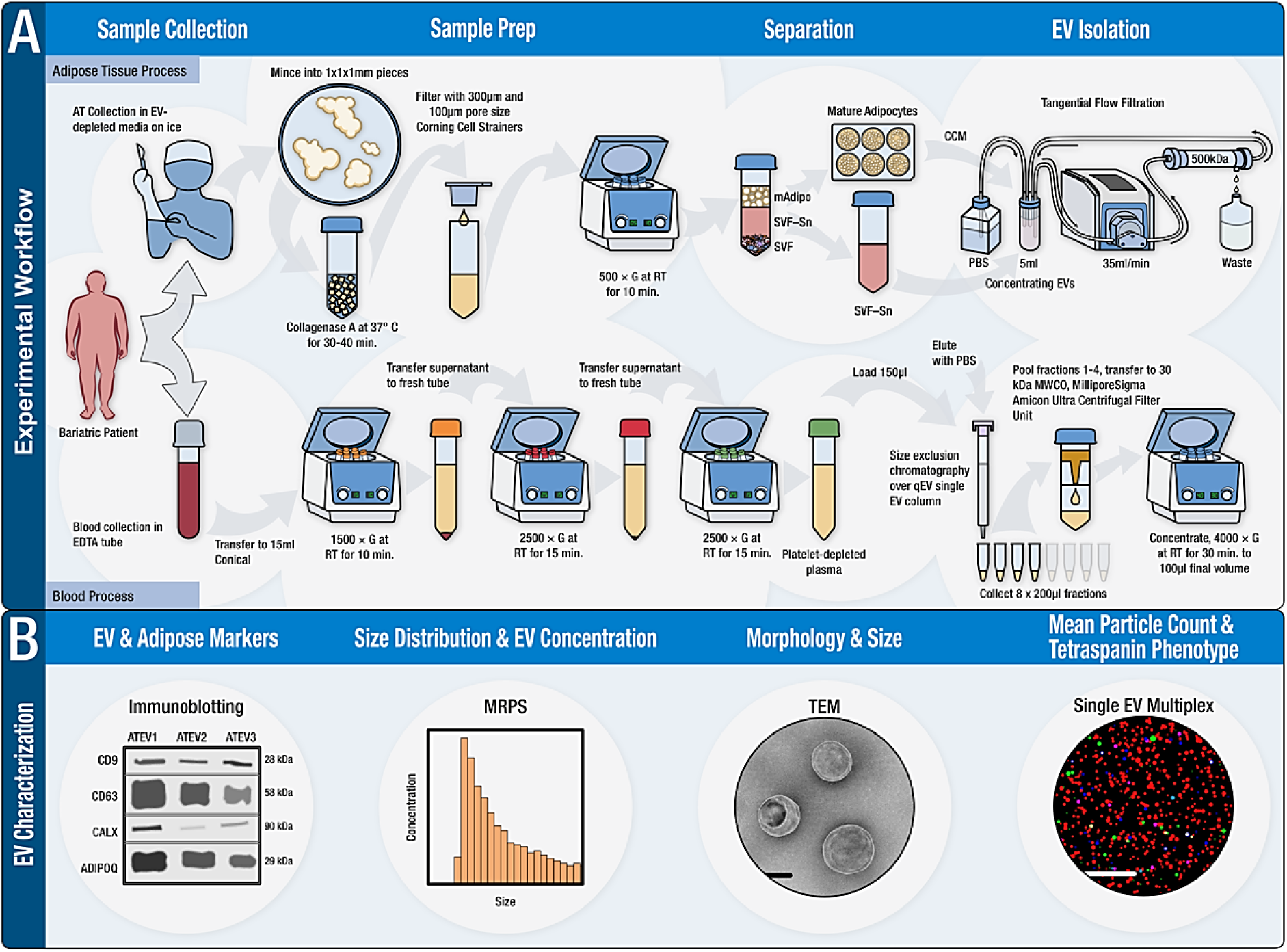
Workflow for EV isolation (A) and characterization (B) of ATEVs, aEVs, and bEVs. **(A)** Visceral adipose tissue (2-10 grams) was enzymatically digested, filtered, and centrifuged, resulting in three phases: pelleted SVF, SVF-SN, and floating mature adipocytes. Mature adipocytes were cultured for 24 hours (see Methods), and the CCM was removed for EV isolation (designated aEVs) via TFF. SVF-SN was also collected for ATEV isolation via TFF. Matched plasma was collected in EDTA tubes and subjected to centrifugation to separate plasma and cellular components. To avoid contamination from platelet EVs, platelet depletion was performed before EV isolation using size exclusion chromatography. Fractions 1-4 were pooled and concentrated. **(B)** Multiparametric characterization of EVs by immunoblotting, MRPS, TEM, and single EV multiplex analysis. ATEVs: adipose tissue EVs, bEVs: blood EVs, SVF: stromal vascular fraction, SVF-SN: stromal vascular fraction supernatant, CCM: cell-conditioned media, TFF: tangential flow filtration, EDTA: ethylenediaminetetraacetic acid, MRPS: microfluidic resistive pulse sensing, TEM: transmission electron microscopy

### Bulk EV characterization of matched human adipose tissue-derived EVs, mature adipocyte EVs, and plasma EVs from the same bariatric patients

Microfluidic resistive pulse sensing (MRPS) was used to analyze the size distribution and particle concentration of EVs from three adipose sources (ATEV, aEV, bEV), all derived from the same individual (**Figure 2A-C**, n = 3 biological replicates). MRPS is an electric sensing platform that utilizes cartridges with pre-specified upper and lower detection limits. Because we aimed to study small EVs, we employed the C400 cartridge, which measures particles within the size range of 65 nm to 400 nm. MRPS results revealed a broader particle size distribution for ATEVs compared to aEVs and bEVs. The median particle diameters were 86.12 nm for ATEVs, 82.26 nm for aEVs, and 78.60 nm for bEVs. Mean particle concentrations were 1.27e+13 particles/mL for ATEVs, 4.09e+12 particles/mL for aEVs, and 4.13e+12 particles/mL for bEVs (**Figure 2A-C**).

**Figure 2.**
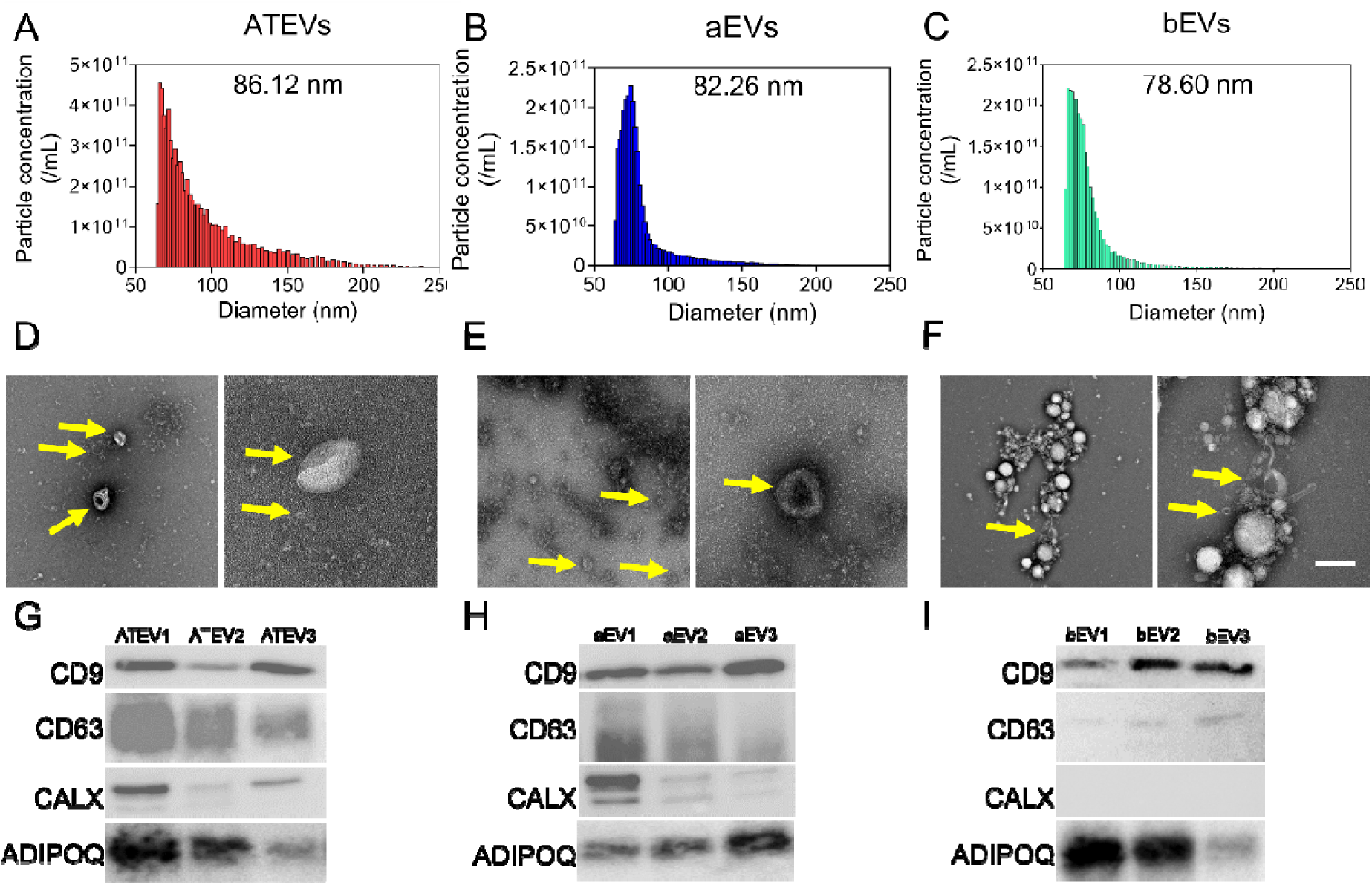
Human adipose EVs exhibit nanoscale size, cup-shaped morphology,and classic EV and adipose markers. **(A-C)** Particle size distribution and concentration analysis via MRPS. **(D-F)** TEM images displaying cup-shaped morphology of isolated EVs. Yellow arrows indicate individual EVs. **(G-I)** Immunoblot analysis of EV (CD9 and CD63), non-EV (CALX), and adipocyte markers (ADIPOQ). N = 3 biological replicates for all analyses. ATEVs: Adipose Tissue-derived Extracellular Vesicles, aEVs: Mature Adipocyte-derived Extracellular Vesicles, bEVs: Plasma Extracellular Vesicles. CD9: Cluster of Differentiation 9, CD63: Cluster of Differentiation 63, CALX: Calnexin, ADIPOQ: Adiponectin. Scale 100 nm.

TEM imaging showed heterogeneous particle populations, including small EVs with classic ‘cup-shaped’ morphology (**Figure 2D-F**). Immunoblot analysis revealed varied protein expression of typical EV/non-EV and adipose markers across biological replicates of ATEVs, aEVs, and bEVs (**Figure 2G-I**). While calnexin is typically considered a non-EV marker, calnexin was detected in ATEVs and aEVs (**Figure 2G**, **H**), consistent with previous reports of endoplasmic reticulum proteins associated with tissue-derived EVs^28–30^. As previously reported, the presence of adiponectin (ADIPOQ) in early SEC fractions suggests its association with EVs rather than free proteins^12^.

### Human ATEVs and aEVs exhibit unique and shared proteomes and biological relevance to obesity-associated metabolic and immune processes

Next, we aimed to develop an ‘adipose EV signature,’ by first performing bulk mass spectrometry-based proteomics (**Figure 3**) to explore the unique and shared proteomes of adipose tissue EVs (reflecting the complex adipose tissue microenvironment) and mature adipocyte EVs (reflecting the ‘purest’ adipose reference material). A three-way Venn diagram revealed 76 unique ATEV proteins, 512 unique aEV proteins, and 1003 shared proteins (**Figure 3A**). The overlapping proteins suggest potential functional similarities and shared signaling pathways, while the unique proteins signify specific molecular characteristics associated with each group. The delineation of shared versus unique proteins provides essential insights into how ATEVs and aEVs contribute differently to biological processes. Additionally, to gain further insights into the protein expression profiles, we performed hierarchical clustering for the top 50 most abundant proteins in ATEVs and aEVs (**Figure 3B**). The distinct clustering observed between ATEVs and aEVs indicates distinct protein expression profiles in each group. The most abundant unique and shared proteins are listed in **Table S2.** This data set was utilized to design the single EV immunocapture probes against adipose-enriched proteins (**Figure 5**) for liquid biopsy development.

**Figure 3.**
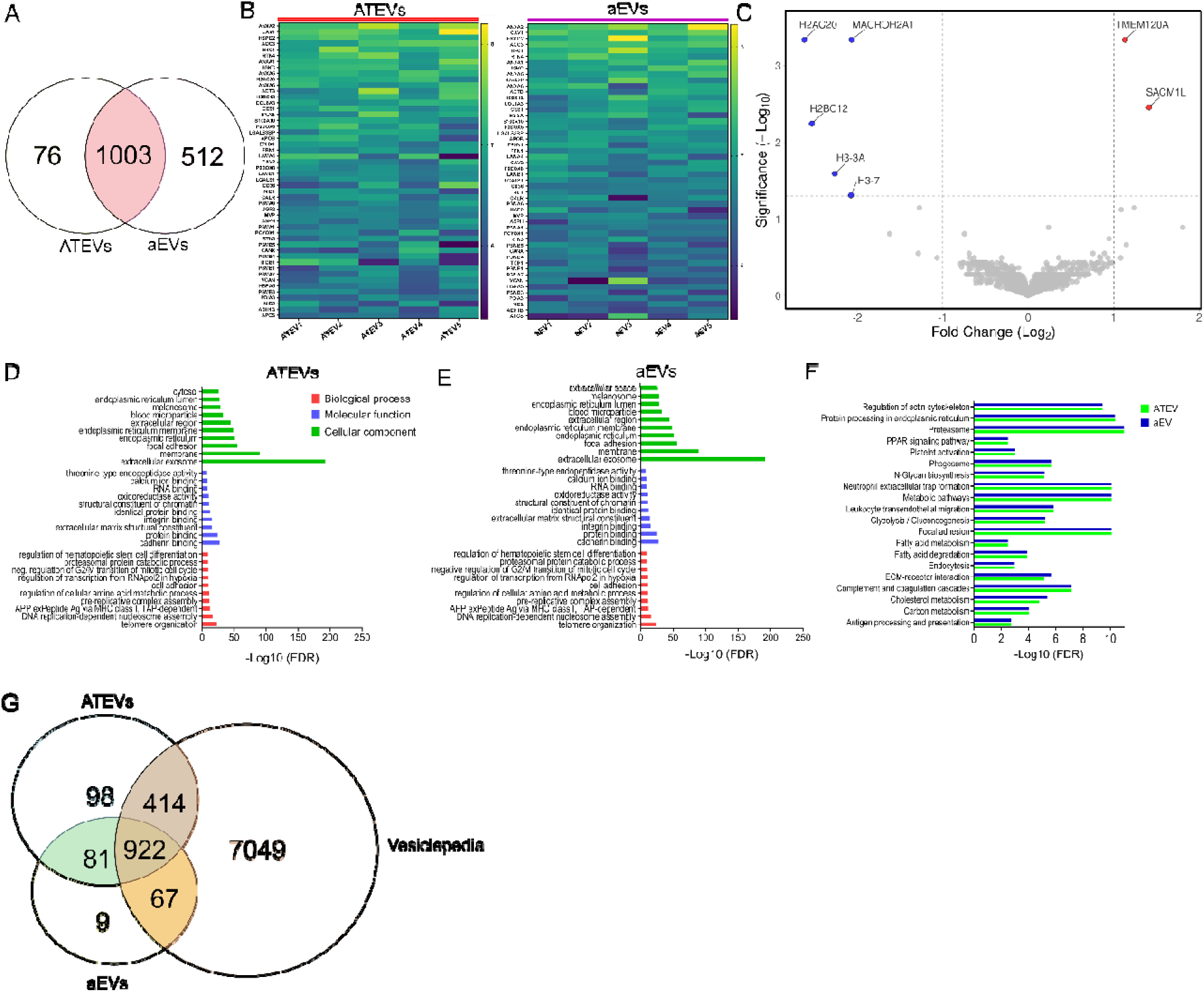
Proteomics analysis of paired human adipose EVs. EVs isolated from matched visceral ATEVs and cultured mature aEVs underwent proteomics analysis. **(A)** A three-way Venn diagram revealed proteins identified in both ATEVs and aEVs (n=1003), with 76 proteins found exclusively in ATEVs and 512 proteins found exclusively in aEVs. **(B)** Heatmaps demonstrated distinct expression profiles of the 50 most abundant proteins from ATEVs (orange) and aEVs (purple), N = 5 biological replicates). **(C)** Volcano plot showed a higher number of upregulated proteins in aEVs (n = 5; H2AC20 (*P* = 0.0004619), MACROH2A1 (*P* = 0.000462), H2BC12 (*P* = 0.0057061), H3-3A (*P* = 0.02572), H3-7(*P* = 0.049126)) compared to ATEVs (n =2; TMEM120A (*P* = 0.000462), SACM1L (*P* = 0.0035054)). Gray points represent proteins that do not show significant differential expression between ATEVs and aEVs. The x-axis represents log2 fold changes (log2(FC)), and the y-axis represents the -log10(p-value). **(D-E)** GO enrichment analysis in ATEVs **(D)** and aEVs **(E)**, including biological process, molecular function, and cellular component subclassifications. **(F)** KEGG pathway analysis of enriched pathways in ATEVs (green) and aEVs (blue). **(G)** Venn diagram demonstrating overlap in protein expression between ATEVs, aEVs and Vesiclepedia EV database. ATEVs: Adipose Tissue-derived Extracellular Vesicles, aEVs: Mature Adipocyte Extracellular Vesicles, GO: Gene Ontology, KEGG: Kyoto Encyclopedia of Genes and Genomes

Volcano plot analysis **(Figure 3C)** revealed significant differences in the proteomic profiles of ATEVs and aEVs. Statistical analysis using the limma method, with thresholds set at an FDR-corrected p-value < 0.05 and a log2 fold change (log2FC) ≥ 1.0, identified two significantly upregulated proteins TMEM120A (*P* = 0.000462), SACM1L (*P* = 0.0035054) in ATEVs. In contrast, five proteins H2AC20 (*P* = 0.0004619), MACROH2A1 (*P* = 0.000462), H2BC12 (*P* = 0.0057061), H3-3A (*P* = 0.02572), and H3-7(*P* = 0.049126) were significantly upregulated in aEVs. Notably, H3-3A and H3-7 showed coordinated upregulation in aEVs, indicating potential involvement in chromatin remodeling or extracellular nucleosome release^31–33^. Similarly, TMEM120A and SACM1L, upregulated in ATEVs, are implicated in lipid metabolism and membrane trafficking^34–37^, respectively, suggesting a possible unique function of ATEVs in these pathways.

To investigate canonical biological processes and molecular functions associated with the identified proteins in both ATEVs and aEVs, we performed Gene Ontology (GO) pathway analysis (**Figure 3D**, **E**). Biological processes identified in our analysis highlighted several significant pathways, including genomic stability, DNA processing, and chromatin remodeling. Our study revealed the involvement of ATEVs and aEVs in processes such as pre-replicative complex assembly, regulation of cellular amino acid metabolic processes, cell adhesion, and regulation of transcription from RNA polymerase II promoter in response to hypoxia. Pathways responsible for antigen processing and presentation, cell adhesion, and protein-protein interactions were also identified.

KEGG pathway analysis provided further insights into the potential pathobiological relevance of ATEVs and aEVs (**Figure 3F**). Prominent and degradation (e.g., proteasome and ER-related pathways), immune responses (e.g., neutrophil extracellular traps, complement cascades, and phagosome formation), cell adhesion and interaction (e.g., focal adhesion and ECM-receptor interaction), metabolism (e.g., glycolysis), and N-glycan biosynthesis. These pathways implicate ATEVs and aEVs in processes related to immune responses, cell adhesion, cellular metabolism, cytoskeletal regulation, and intercellular communication. Moreover, pathways associated with cholesterol metabolism, carbon metabolism, fatty acid degradation, platelet activation, endocytosis, antigen processing and presentation, PPAR signaling, and fatty acid metabolism were also identified.

To provide additional context, we compared our ATEVs and aEVs data to Vesiclepedia^38^ (**Figure 3G**), an open-access database that compiles experimentally identified proteins from various EV subtypes. A three-way Venn diagram revealed 922 shared proteins among ATEVs, aEVs, and Vesiclepedia, representing conserved EV markers. Vesiclepedia contributes 7049 unique proteins, while ATEVs and aEVs exhibit 98 and 9 unique proteins, respectively. Pairwise comparison revealed 414 proteins shared between ATEVs and Vesiclepedia, 67 proteins between aEVs and Vesiclepedia, and 81 proteins between ATEVs and aEVs. These results highlight both unique and shared contributions across EV dataset Vesiclepedia^38^.

In summary, the combined proteomic and functional analyses revealed distinct molecular signatures in ATEVs and aEVs. ATEVs are involved in cellular structure, adhesion, and communication, reflecting their role in maintaining the integrity of adipose tissue. In contrast, aEVs play a more specialized role in metabolism and immune-related processes, indicating their involvement in protein handling and immune modulation. These findings provide insights into the unique and shared biological roles of ATEVs and aEVs and possible contributions to obesity-related metabolic and immune processes.

### Distinct protein interaction networks and functional insights of ATEV and aEV proteins

To gain insights into the role(s) of proteins from aEVs and ATEVs, we generated protein-protein interaction (PPI) networks, highlighting the most enriched proteins **(Table S2)** and their interacting partners (**Figures 4** **and S3-S22, Table S3-S10**). Several proteins were shared between ATEVs and aEVs (**Figure 4**), such as ANXA2 (Annexin A2), CAV1 (Caveolin-1) and RTN3 (Reticulon-3). ANXA2 interacts with S100A10 and PLIN1, facilitating membrane repair, trafficking, and vesicle fusion, which are crucial during metabolic stress and tissue remodeling (**Figure 4A**). CAV1 supports lipid raft formation, cholesterol homeostasis, and vesicular transport, interacting with membrane-associated proteins to regulate endocytosis and intracellular signaling ^39–41^ (**Figure 4A**, **B**). RTN3 participates in membrane curvature and vesicle trafficking, helping maintain membrane integrity during stress conditions^42–44^ (**Figure 4C**, **S7**).

**Figure 4.**
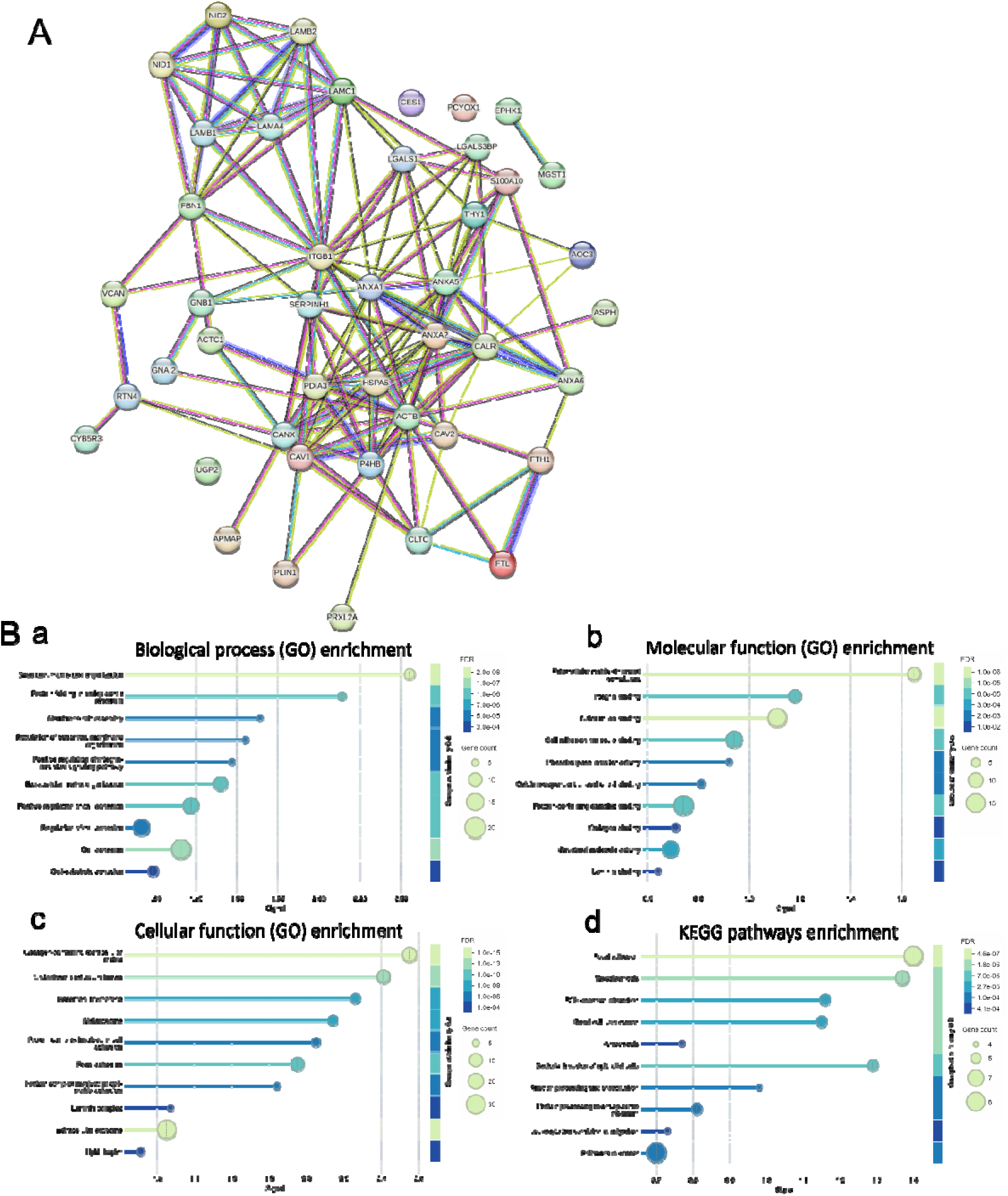
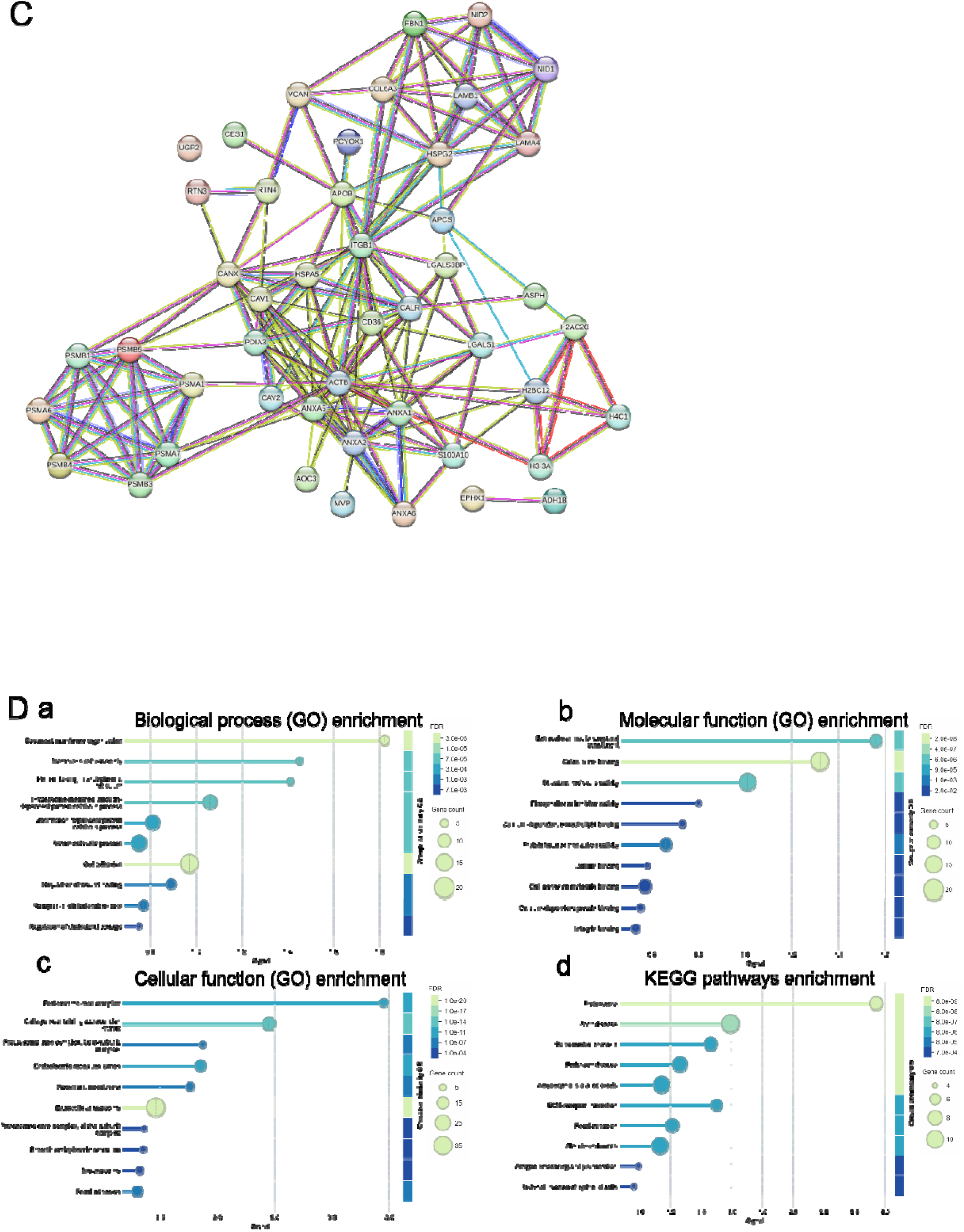
Interaction map for the top 50 proteins identified in ATEVs and aEVs. **(A)** Protein protein interaction networks for the top 50 ATEV proteins **(B)** ATEV protein functional enrichment GO analyses for (a) biological process, (b) molecular function, (c) cellular function, and (d) KEGG pathway enrichment. **(C)** aEV protein-protein interaction networks for the top 50 proteins. **(D)** Go enrichment analyses for (a) biological process, (b) molecular function, (c) cellular function, and (d) KEGG pathway enrichment. Protein interaction networks were generated using STRING database integrated with Cytoscape software (v3.10.2) with a minimum interaction score threshold of 0.7 for significantly differentially expressed proteins.

ATEV network proteins were involved in systemic metabolic regulation and immune modulation (**Figure 4A**, **B**). LAMC1 (Laminin Subunit Gamma-1) (**Figure S3**) and LAMB2 (Laminin Subunit Beta-2) (**Figure S4**) interact with extracellular matrix components, supporting cell adhesion, matrix organization, and tissue integrity^45–48^. These interactions are essential for interorgan communication and structural maintenance in metabolic tissues ^45,48^. Cytochrome B5 Reductase 3 (CYB5R3) regulates redox balance and lipid metabolism by interacting with enzymes involved in fatty acid desaturation and steroid biosynthesis^49–51^ (**Figure S5**). PRXL2A (Peroxiredoxin-Like 2A) supports antioxidant defense, interacting with stress-response proteins to mitigate oxidative damage during inflammation^52,53^ (**Figure S6**). RTN3 (Reticulon 3) was found to be a central node interacting with key proteins involved in vesicle trafficking, endoplasmic reticulum (ER) organization, and autophagy. Notably, RTN3 interacts with RTN4 and REEP5, which support ER shaping and intracellular trafficking^54^ (**Figure S7**). RTN3 also interacts with autophagy-related proteins like GABARAP and GABARAPL2, emphasizing its role in autophagosome formation^55,56^. Moreover, interactions with BACE1 and BACE2 were also identified and suggest involvement in amyloid precursor protein processing, linking this network to neurodegenerative pathways^57^ (**Figure S7**).

Additional ATEVs networks included FABP4 (Fatty Acid Binding Protein 4), which interacts with ACSL1 (Acyl-CoA Synthetase Long Chain Family Member 1) to promote fatty acid transport and utilization ^58–60^. GNAI2 (G-protein subunit alpha-2) regulates hormonal signaling pathways that influence energy metabolism and insulin sensitivity^61^ (**Figure S8, Table S2**). FTL (Ferritin Light Chain) interacts with iron-binding proteins to regulate iron homeostasis. This interaction is crucial in managing oxidative stress since excess iron contributes to the generation of reactive oxygen species (ROS), further linking ATEVs to systemic immune regulation^62,63^ (**Figure 4A**, **B**, **S9, Table S2**). CLTC (Clathrin Heavy Chain) interacts with adaptor proteins AP2A1, AP2B1, and AP2M1 to regulate vesicle formation and cargo sorting, while partners like HIP1 and GAK link endocytosis with cytoskeletal dynamics for efficient membrane recycling ^64,65^ (**Figure 4A, B, S10, Table S2**). Immune-related proteins such as C3 (Complement C3) interact with C5, contributing to immune surveillance and inflammation control, which is critical in obesity-related metabolic dysfunction^66^. The ATF1 interaction network (**Figure S11**) with partners such as CREB1, ATF2, and CREB3L, highlights their collective roles in transcriptional regulation^67^, chromatin binding^68^, and key metabolic pathways like insulin and TNF signaling^69^. The interaction network of ACTC1(**Figure S12**) with partners such as TPM1, TNNT2, MYH6, and MYL2, emphasizes their roles in muscle contraction, cytoskeletal organization, and cardiac function^70^.

In contrast, the aEV network focuses on local tissue remodeling, chromatin regulation, and oxidative stress management (**Figure 4C, D**). HSPG2 (Perlecan) interacts with matrix proteins to support angiogenesis, tissue elasticity, and extracellular matrix stability, which are essential during adipogenesis and tissue expansion^71–73^ (**Figure 4C, D, S13, Table S2**). Several histones, including H4C1, H2AC20, H2BC12, and H3-3A, are enriched in aEVs, indicating their involvement in chromatin remodeling and gene expression regulation^31,33,74^(**Figure 4C, D, S14, S15, Table S2**). These interactions ensure that adipocytes efficiently regulate transcription to accommodate lipid storage and cellular differentiation^32,75,76^. Additionally, MGST1 (Microsomal Glutathione S-Transferase 1) interacts with glutathione pathways, contributing to detoxification and oxidative stress management within the adipose microenvironment^77,78^ (**Figure S16, Table S2**). PSMB5 (Proteasome Subunit Beta Type-5) interacts with other proteasome core components such as PSMB6, PSMB7, and PSMB1 to form the 20S core of the proteasome, essential for protein degradation (**Figure 4C, D, S17, Table S2**). Additionally, it interacts with PSMA3, PSMA4, and PSMA6, facilitating the assembly of the proteasome’s catalytic core, which is critical for regulating cellular protein turnover, stress responses, and maintaining protein homeostasis^79^ (**Figure 4C, D, S17, S18, Table S2**). H4C6 (Histone H4 Cluster Member 6) interacts with several histone proteins, including H3-3B, H2AC20, and H3C12, forming part of the nucleosome core responsible for chromatin structure and regulation^80^ (**Figure 4C, D, S19**). It also interacts with BRD4 and EP300, proteins involved in histone acetylation and transcriptional activation, highlighting its role in facilitating gene expression and epigenetic regulation during adipocyte differentiation^81,82^ (**Figure S19**). APOB (Apolipoprotein B) interacts with MTTP, APOE, and SCARB1 to mediate lipoprotein assembly, lipid transport, and cholesterol uptake, playing a crucial role in lipid homeostasis and cardiovascular health^83,84^ (**Figure S20, Table S2**). The APCS (Serum Amyloid P-Component) network highlights interactions with CRP, TTR, and APOE, indicating a role in immune response and lipid transport^85^. Additional interactions with FN1 and FCGR2A suggest involvement in extracellular matrix organization and receptor-mediated immune functions^86^ (**Figure 4C, D, S21, Table S2**). CD36 interacts with TLR2, TLR4, and TLR6, linking lipid metabolism with innate immune signaling and inflammatory responses (**Figure S22, Table S2**)^87^. CES1 (Carboxylesterase 1) interacts with enzymes like CYP3A4, UGT1A6, and NCEH1, playing a key role in lipid metabolism, detoxification, and hydrolysis of esters and triglycerides^88^ (**Figure S23, Table S2**).

The presence of both ANXA2 (**Figure 4**) and PRXL2A (**Figure 4**) in aEVs and ATEVs highlights their shared role in oxidative stress management^89,90^. However, the functional emphasis diverges: aEVs primarily maintain adipose tissue structure, supporting local tissue remodeling and plasticity, while ATEVs function as systemic endocrine messengers, coordinating metabolic responses, immune signaling, and lipid metabolism across tissues.

These findings suggest that targeting specific EV populations could offer novel therapeutic opportunities. Modulating ATEVs may enhance systemic metabolic health and manage chronic inflammation, while enhancing aEV content could optimize tissue remodeling and improve insulin sensitivity. This functional divergence highlights the potential of aEVs and ATEVs as biomarkers and therapeutic targets for metabolic diseases.

Edges indicate protein-protein interactions, represented by different colors: Known Interactions (Curated Databases: Light blue line, Experimentally Determined: Pink line), Predicted Interactions (Gene Neighborhood: Green line, Gene Fusions: Red line), Gene Co-occurrence: Blue line, Others (Text mining: Yellow line, Co-expression: Black line, Protein Homology: Lavender line). See supplemental Tables S11 and S12 for detailed protein descriptions.

### Multiparametric single EV analysis of human adipose-derived EVs

Orthogonal and complementary single EV methods were utilized to characterize EV size, tetraspanin distribution, EV and adipose protein targets, and select adipose mRNA transcripts in paired ATEVs and aEVs from the same individual.

### Single ADEV analysis leveraging single-particle interferometric reflectance imaging sensor (SP-IRIS)

SP-IRIS^91^ with fluorescence was employed to gain insights into the complexity and heterogeneity of tetraspanin distribution across biologically similar but distinct adipose-enriched EV subsets at single EV resolution (**Figure 5**). Single EVs were immunologically immobilized/captured by surface tetraspanin proteins CD63, CD81, and CD9 printed on a microchip surface. Fluorescently-labelled anti-CD63, anti-CD81 and anti-CD9 antibodies were added for detection. Capture events were counted (mean particle count) and sEVs were sized by light interference (lower limit of detection of 50 nm). SP-IRIS revealed a particle size scatter of 50 nm to 200 nm for both ATEVs and aEVs (data not shown), which was consistent with MRPS size analysis. Quantification of events per capture chip revealed a notable presence of particles, indicating successful ATEV (**Figure 5A, C**) and aEV (**Figure 5B, D**) capture. Both ATEVs and aEVs demonstrated the most total capture events with CD63 capture and greater CD63 detection from the CD63 capture spot. CD81 capture resulted in similar expression patterns for ATEVs and aEVs while CD9 capture revealed higher CD9 detection for aEVs but not ATEVs (**Figure 5A, B**).

**Figure 5.**
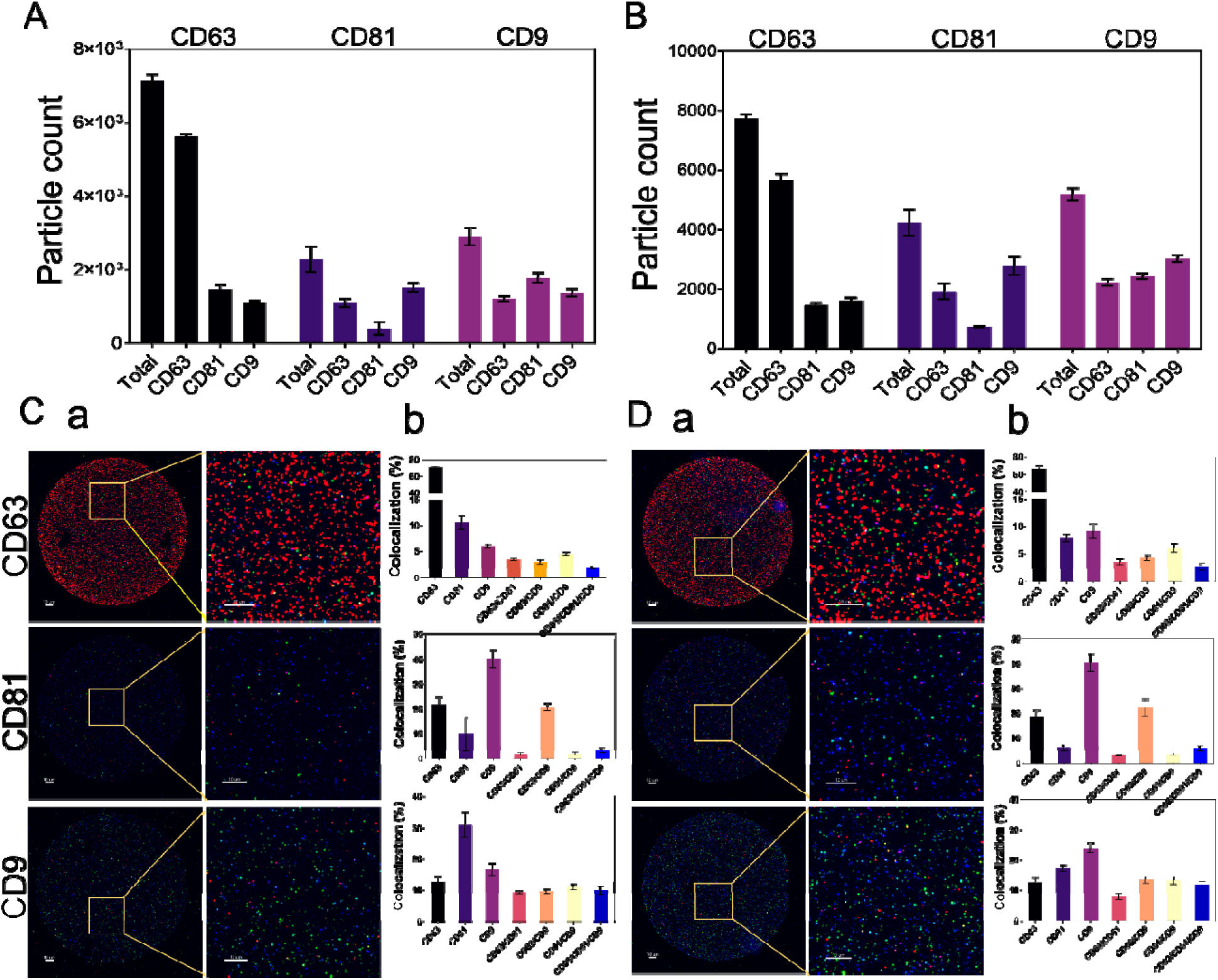
Single ADEV multiplex analysis showed comparable classic EV markers in ATEVs and aEVs. Tetraspanin profile characterization for ATEVs (**A**) and aEVs (**B**). The x-axis reflects the tetraspanin detection antibody and the top label indicates the tetraspanin capture antibody. Both ATEVs and aEVs showed similar but not identical tetraspanin event frequencies (see manuscript text for details; n =3 biological replicates). SP-IRIS representative images per capture spot CD63, CD81, CD9 for ATEVs (**Ca**) and aEVs (**Da**). Scale 10 um. All events were detected above mouse IgG isotype control levels (data not shown). Tetraspanin colocalization analysis for ATEV (Cb) and aEV (Db). Fluorescently-labeled anti-CD63, anti-CD81, and anti-CD9 were used for particle detection after particle capture with CD63, CD81, and CD9. Capture antibody is indicated in the upper right corner of each graph. In ATEVs, CD9 and CD81 show similar co-expression levels. In aEVs, CD9 positive particles demonstrate less degree of co-expression with CD81.

Tetraspanin colocalization analysis revealed distinct patterns among the CD63, CD9, and CD81 capture spots **(Figure 5C, D; Figure S1)**. CD63 was the most prevalent single-positive tetraspanin detected on both ATEVs and aEVs immobilized on the CD63 capture spot, as visualized by the red fluorescence signal (**Figure 5C, D**). ATEVs and aEVs captured by CD81 both had the highest detection for CD9+ single-positive EVs, as well as CD63+/CD9+ double-positive EVs. CD9 capture differed between ATEVs and aEVs, with greater CD81+ single positive ATEVs and slightly greater CD9+ single positive aEVs. These results indicate that CD63 is the predominant surface marker expressed in both ATEVs and aEVs. Representative images obtained with the SP-IRIS instrument visually confirmed these expression patterns (**Figure 5C, D**). In summary, Single EVs tetraspanin phenotyping provides valuable insights into the tetraspanin expression profiles and molecular heterogeneity of ATEVs and aEVs.

### Total internal reflection fluorescence (TIRF) microscopy allows simultaneous detection of human adipose-derived EV protein and mRNA cargo at single EV resolution

To establish an adiposity-associated EV liquid biopsy for the interrogation of circulating plasma EVs, we next applied our adipose-enriched proteomics candidates to develop adipose-enriched surface capture targets and molecular beacons for mRNA detection cargo at single EV resolution. After immobilization of ATEVs and aEVs for proof of principle by using an anti-CD63 and CD9 hybridized surface^92^, two adipocyte markers (adipocyte plasma membrane protein, APMAP; and perilipin 1)^93,94^ were selected for detection of adipocyte-specific surface proteins on plasma EVs. Furthermore, we examined the presence of specific mRNA molecules using molecular beacons designed against PPARG and ADIPOQ, which are known to be highly expressed in adipocytes^95–96^. The results of the protein detection revealed the presence of APMAP, perilipin, and CD63 on both single ATEVs and aEVs (**Figure 6**). Similarly, detection of the molecular beacons confirmed the presence of PPARG and ADIPOQ mRNA within the immobilized vesicles. These results confirm our ability to characterize plasma circulating, adipose-specific single EVs with simultaneous surface protein and mRNA cargo.

**Figure 6.**
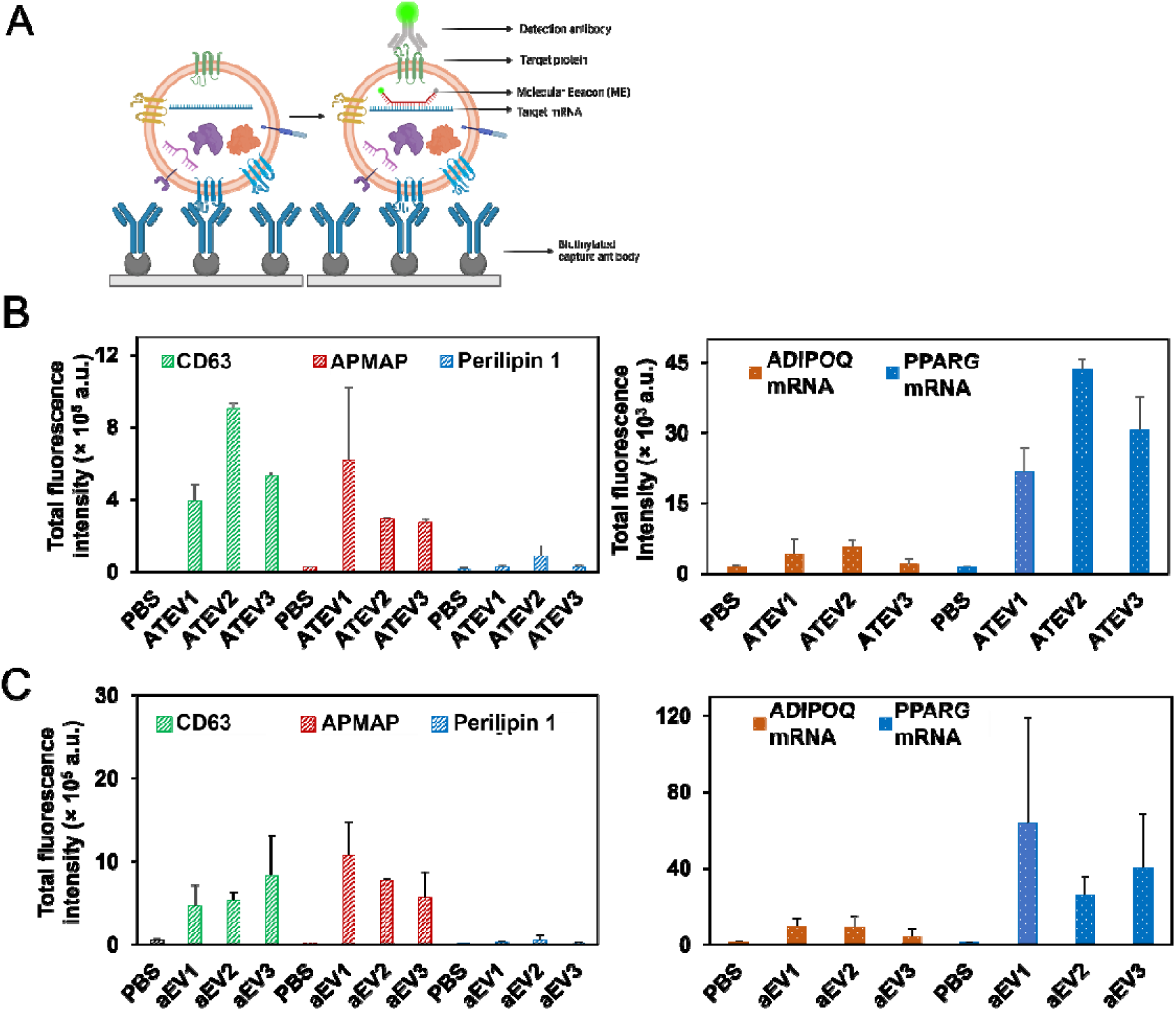
Single ADEV cargo and surface marker analysis defines CD63+/APMAP+/PPARG+ as an adipose-specific EV signature for liquid biopsy development. **(A)** A schematic illustration depicting the working principle, demonstrating the operational mechanism of TIRF. **(B)** ATEV and **(C)** aEV TIRF analysis with simultaneous protein (APMAP, perilipin 1, CD63; left) and mRNA cargo (ADIPOQ, PPARG; right) (n = 3 paired biological replicates). ADIPOQ: Adiponectin, APMAP: adipocyte plasma membrane-associated protein, PPARG: peroxisome proliferator-activated receptor gamma, ADEV: adipose-derived EVs, ATEVs: adipose tissue-derived EVs, aEVs: mature adipocyte-derived EVs

### Multiparametric single particle analysis of plasma EVs and validation of an adiposity-associated EV liquid biopsy

Lastly, to explore the liquid biopsy utility of our adipose-derived EV analysis, we applied two complementary single EV methods to the plasma EVs (bEVs) that corresponded to the matched ATEVs and aEVs from the same individual (**Figure 1,5,6**). SP-IRIS analysis was used to explore the tetraspanin distribution of bEVs. Unlike the matched ATEV and aEV analyses, CD81 capture, rather than CD63 capture, resulted in the greatest total mean particle count and CD81+ single positive EVs (**Figure 7A, B**). Colocalization analysis demonstrated a high percentage of CD63+/CD81+ double-positive bEVs (**Figure 7Bb, Figure S2**). These results underscore the heterogeneity of EV subsets across various biospecimens.

**Figure 7.**
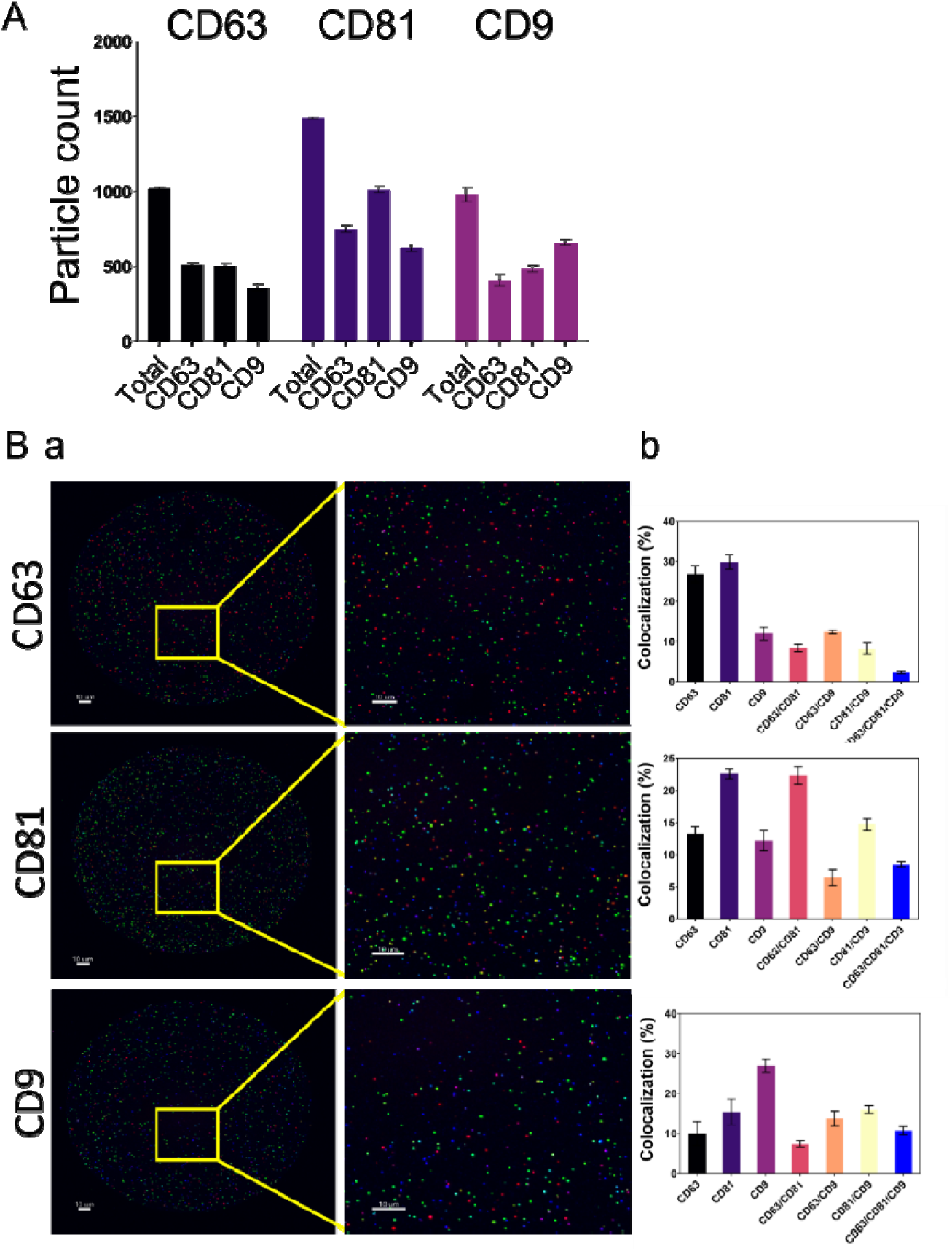
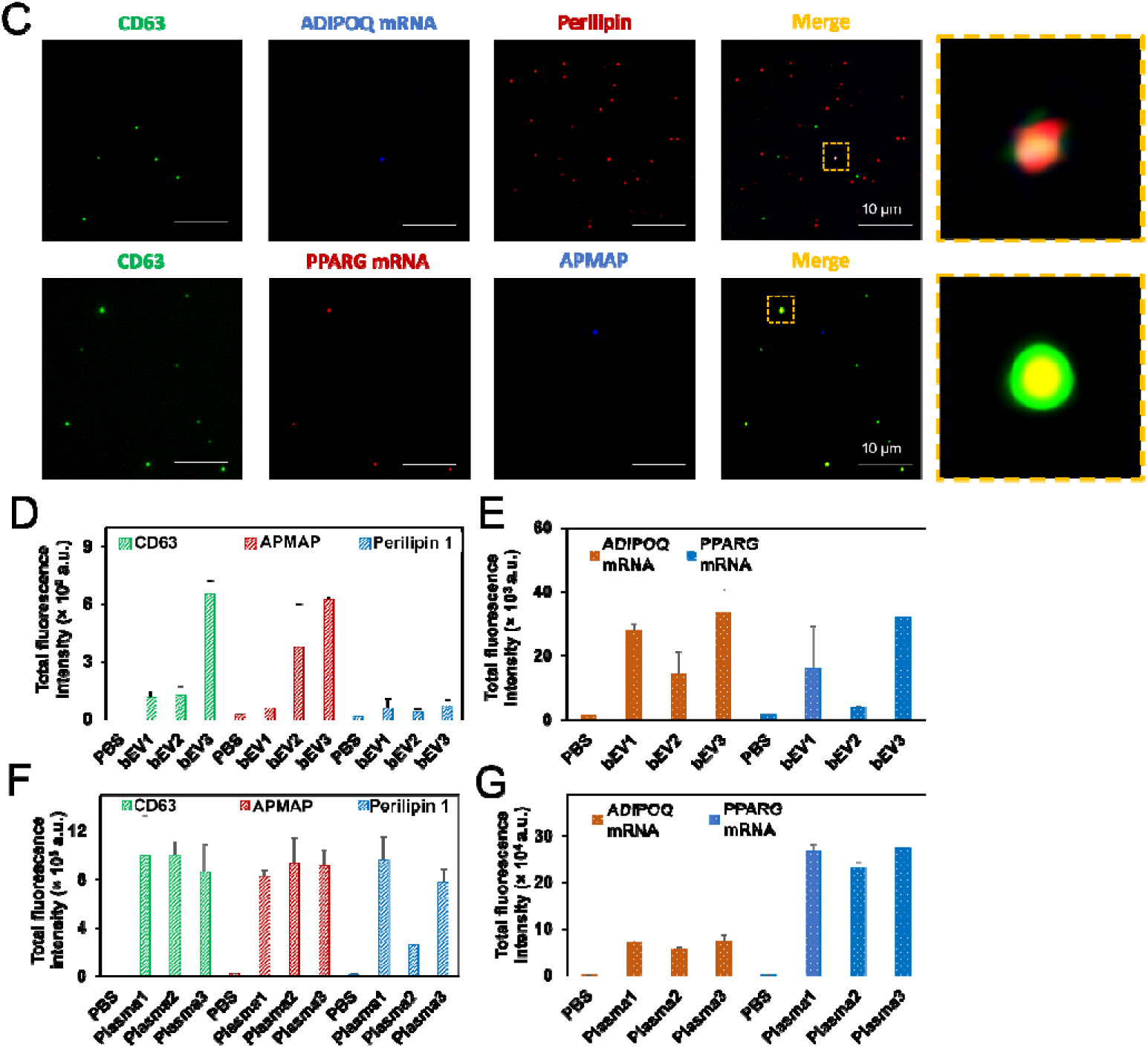
Multiparametric single EV analysis of plasma EVs with simultaneous adipose target protein and mRNA cargo characterization. **(A, B)** SP-IRIS tetraspanin profile characterization and colocalization representative images of bEVs (scale 10 µm). Tetraspanin colocalization analysis for plasma EVs (bEVs) **(B) (a)** Fluorescently-labelled anti-CD63, anti-CD81, anti-CD9 were used for particle detection after particle capture with CD63, CD81, and CD9. (b) Colocalization analysis demonstrated a high percentage of CD63+/CD81+ double-positive bEVs. **(C)** TIRFM images showing fluorescent colocalization of proteins and mRNAs on single bEVs. Insets show enlarged images of single bEVs. Total fluorescence intensity analysis on paired SEC-isolated plasma EVs **(D-E**) and neat plasma (**F-G**) from the same individual. Simultaneous adipocyte and EV protein (APMAP, perilipin 1, CD63; left) and adipocyte mRNA cargo (ADIPOQ, PPARG; right) detection in single bEVs from three separate bariatric patients (n = 3 matched biological replicates).

TIRF analysis of bEVs (**Figure 7C-E**) and neat plasma (**Figure 7C, F, G**) was employed to validate our ability to capture and characterize adipose-derived EVs in the circulation of bariatric patients. We successfully captured isolated, plasma-derived EVs that displayed EV (CD63) and adipocyte surface markers (APMAP, perilipin 1) and contained adipocyte mRNA cargo (ADIPOQ, PPARG) (**Figure 7D-E**). We next sought to evaluate the sensitivity of our bEV assay to detect low-abundant ADEVs in the circulation of bariatric patients by performing TIRF analysis on neat plasma (i.e., without EV isolation) from the same patients. We were again able to capture bEVs that displayed EV (CD63) and adipocyte surface markers (APMAP, perilipin 1) and contained adipocyte mRNA cargo (ADIPOQ, PPARG) (**Figure 7F, G**).

## Discussion

The dual roles of ADEVs as both biomarkers and pathogenic conveyors of disease across a diversity of pathologies have recently garnered increased attention^4,21,22,97^. Through the transfer of diverse bioactive cargoes, ADEVs can influence disease mechanisms locally and systemically^22,98–100^, particularly at the interface between adipose tissue and systemic metabolic regulation^101^. Despite the growing interest in ADEVs, their full translational potential has been hindered by methodological and biological challenges. Firstly, standardized, reproducible and robust isolation protocols for tissue-derived EVs are lacking^25^. Secondly, most ADEV studies have focused on rodent models^21–23^, despite known cross-species differences in adipose biology between rodents and humans^24^. Lastly, cell-specific EV analysis is often technically challenging and also requires a priori knowledge of known cell-specific markers for EV immunocapture^17^. Therefore, innovative approaches for human adipocyte-specific EV analysis are needed to move the translational potential of ADEVs beyond the laboratory and into the clinic.

This study establishes a robust, multiparametric methodological framework for isolating and characterizing ADEVs using a combination of bulk and single-EV analytical techniques. By integrating advanced proteomics and single-EV approaches, including TIRF microscopy, we address existing challenges in ADEV research, such as reproducibility, specificity, and sensitivity in identifying tissue-derived EV markers.

### A Multiparametric framework for ADEV characterization

Our approach combines bulk proteomics with cutting-edge single-EV techniques to provide a detailed understanding of ADEV heterogeneity. By leveraging paired human biospecimens including VAT, mature adipocyte-conditioned media, and plasma, we ensured comprehensive coverage of ADEVs across tissue, cell, and systemic sources. This pipeline establishes the feasibility of characterizing low-abundance ADEVs in biofluids, which has historically been a significant technical challenge due to the complexity of plasma EVs^10^. One of the key strengths of this framework is its ability to consistently capture intra-individual similarities across ADEV sources, despite inter-individual variability.

This finding underscores the reliability of ADEVs as biomarkers for obesity and related diseases, including cardiovascular disorders, autoimmunity, and neurodegenerative diseases. Importantly, it also highlights the translational potential of ADEVs in clinical diagnostics and liquid biopsy applications.

Our proteomic analysis highlights distinct and overlapping protein signatures in ATEVs and aEVs, shedding light on their complementary roles in adipose tissue biology. ATEVs were enriched with proteins involved in extracellular matrix organization^102^, lipid metabolism^103,104^, and immune modulation^104,105^, while aEVs demonstrated specificity for chromatin remodeling^104^ and oxidative stress responses^106,107^. Importantly, we validated key proteomic findings at the single-EV level using TIRF microscopy, which allowed simultaneous detection of adipose-enriched protein (APMAP and perilipin) and mRNA cargo (PPARG and ADIPOQ). This linkage between proteomic data and single-EV characterization establishes the reliability of our workflow and demonstrates the feasibility of using these markers for ADEVs.

The application of TIRF microscopy in this study proved instrumental in confirming the presence of adipose-specific proteins and transcripts within ADEVs. By capturing EVs using EV specific markers like CD63 and CD9, we demonstrated the co-detection of ADEV proteins and mRNAs associated with adipocyte-specific functions. These findings not only validated our proteomic results but also underscored the utility of TIRF microscopy for detecting low-abundance, tissue-specific EV subsets. This approach has broader implications, as it can be adapted to investigate other protein types and cargo in diverse EV populations, thereby extending its applicability to other studies. TIRF has the advantage of being simpler and less expensive than the alternative immunohistochemistry MALDI mass spectrometry (MS) imaging approach that might also be used, while the MS approach will likely allow higher level multiplexing.^108^

Moreover, our methodology demonstrates how single-EV analysis can complement bulk characterization. Proteomic data provided a comprehensive view of ADEV protein content, while TIRF microscopy enabled high-resolution validation at the single-vesicle level. Together, these methods establish a pipeline for the sensitive and specific analysis of ADEVs, which can be applied to other EV populations for biomarker discovery or therapeutic validation.

This study paves the way for using ADEVs as biomarkers in obesity and related disorders such as cardiovascular disease^2^, cancer^109^, autoimmunity^110^ and neurodegenerative diseases^6^. By developing a framework that identifies and validates adipose-specific EV markers across biospecimens, we contribute to the potential utility of ADEVs in liquid biopsies. The integration of proteomics and single-EV technologies enables precise characterization, enhancing the translational value of EV research. Future studies can leverage this pipeline to explore additional EV populations and their roles in various pathophysiological contexts.

### Biological insights from adipose-derived EV proteomics

Adipose-derived EVs (such as ATEVs and aEVs) are central players in the growing field of adipose tissue interorgan crosstalk and provide added complexity and scope to the adipose secretome^100,101,111^. Our proteomic analysis highlights distinct and overlapping protein profiles between ATEVs and aEVs, emphasizing their complementary roles in metabolic regulation and immune modulation. A total of 1003 proteins were shared between ATEVs and aEVs, while 76 and 512 proteins were uniquely expressed in ATEVs and aEVs, respectively. These findings reflect both the heterogeneity and specialization of these EV populations, offering new insights into how different adipose-derived vesicles contribute to obesity-associated processes.

### Unique molecular features of ATEVs and aEVs

The unique enrichment of proteins involved in cytoskeletal regulation, extracellular matrix (ECM) organization, and immune signaling pathways in ATEVs reflects their critical role in maintaining tissue architecture and modulating inter-organ communication. Key proteins identified in ATEVs include LAMC1 and LAMB2, which are involved in matrix remodeling and cell adhesion, supporting structural integrity in metabolic tissues^45–48^. Additionally, the presence of FABP4 and CLTC^64,65^ in ATEVs highlights their involvement in lipid transport and immune responses, further emphasizing their systemic metabolic relevance^58–60^.

Conversely, aEVs, which represent a more specialized subset, are enriched in histone proteins (H2BC12, H2AC20, H3-3A), suggesting that these EVs are involved in chromatin remodeling and transcriptional regulation^31,33,74^. This specialization aligns with the role of aEVs in metabolic adaptation and oxidative stress responses within adipocytes, which are critical for managing lipid storage and cellular stress. The differential protein expression profiles identified between ATEVs and aEVs were visualized through heatmaps and volcano plots, confirming distinct functional roles across these EV subsets.

### Proteasomal and autophagosomal regulation in aEVs and ATEVs

Adipocyte EVs and ATEVs exhibited a notable enrichment in proteins involved in proteasomal degradation (PSMD12, PSMD3)^112–114^ and autophagosomal regulation (SACM1L)^115,116^, reflecting their involvement in maintaining intracellular protein homeostasis. Proteins like SACM1L, a phosphatidylinositol-4-phosphate-specific phosphatase, are essential for autophagy and intracellular trafficking^115,117^, underscoring the contribution of ADEVs to stress responses and systemic metabolic regulation. These findings indicate that ADEVs support adipose tissue function by facilitating the degradation of misfolded proteins and regulating cellular responses to metabolic stress.

### Lipid metabolism and immune modulation in ATEVs and aEVs

The proteomic analysis also identified key lipid metabolism regulators across both ATEVs and aEVs. FABP4 and ACSL1 were predominantly found in ATEVs, promoting fatty acid transport and utilization, which are essential processes for systemic energy regulation^59,60,118^. The presence of CAV1 in ATEVs further highlights their role in lipid raft formation and cholesterol homeostasis, essential for metabolic signaling pathways^39–41^. These functions align with the broader role of ATEVs in inter-organ metabolic communication and immune modulation, especially in obesity, where systemic inflammation is prevalent.

In contrast, aEVs were enriched in oxidative stress management proteins, including MGST1 (Microsomal Glutathione S-Transferase 1). This protein supports detoxification processes and helps mitigate oxidative damage in adipocytes, a key challenge in obesity-driven metabolic dysfunction^77,78^. These findings emphasize that while ATEVs contribute to systemic metabolic regulation, aEVs play a more localized role, protecting adipocytes from oxidative stress and ensuring proper metabolic function within adipose tissue.

### Shared networks and pathway analysis

Despite their distinct functions, ATEVs and aEVs also share several important proteins that facilitate vesicle trafficking and intracellular communication. Proteins such as ANXA2^89,90^ and RTN4^42–44^ were found in both EV types, contributing to membrane repair, vesicle curvature, and ER organization. These shared molecular features suggest that both ATEVs and aEVs work together to maintain tissue homeostasis and coordinate metabolic and immune responses.

Pathway enrichment analysis further revealed that ATEVs are involved in immune modulation, protein degradation, and ECM organization, while aEVs play a critical role in glycolysis, gluconeogenesis, and oxidative stress management. These distinct pathway enrichments align with the complementary roles of these EV types in obesity: ATEVs support immune regulation and tissue integrity, whereas aEVs focus on metabolic regulation and stress adaptation.

### Clinical relevance

The distinct molecular profiles of ATEVs and aEVs have important implications for obesity-related diseases, including cardiovascular disease, insulin resistance, and chronic inflammation. ATEVs, with their enrichment in ECM-related proteins, may contribute to fibrosis and atherosclerosis, while aEVs, with their focus on oxidative stress responses, may help mitigate insulin resistance and metabolic dysregulation. These findings highlight the therapeutic potential of targeting specific EV populations to improve metabolic health and reduce obesity-related complications.

### Single ADEV multiplex analysis as a next-generation tool for adiposity liquid biopsy development: Method validation through single-EV analysis

This study also leveraged two complementary, next-generation microchip-based platforms to comprehensively phenotype paired human ADEVs from multiple sources at single EV resolution^17^. SP-IRIS^91^ and TIRF microscopy have emerged as valuable tools for the visualization, quantification, and characterization of individual EVs^119,120^. By combining these complementary techniques, we addressed critical challenges in EV research, including the molecular heterogeneity of EV populations^91^ and their relevance to liquid biopsy applications. SP-IRIS analysis enabled single-EV tetraspanin profiling, revealing that while CD9 and CD63 are widely considered pan-EV markers, their expression profiles are not uniform across EV subsets. Consistent with prior reports, our findings demonstrate that tetraspanin expression is highly heterogeneous, reflecting the cellular origin of the EVs. Notably, we found CD63 to be the predominant tetraspanin in ATEVs and aEVs but not in plasma EVs from the same individuals. This observation underscores the importance of assessing multiple surface markers for comprehensive EV phenotyping and highlights the need to consider source-specific tetraspanin variability when interpreting EV data. These results highlight the molecular heterogeneity of EV subsets from different biospecimens and underscore the need to use multiple EV surface markers for immunocapture and/or characterization^91^.

To complement these insights, we utilized TIRF microscopy, a high-resolution imaging technique capable of simultaneously detecting proteins and mRNA molecules within immobilized paired ATEVs, aEVs, bEVs and neat plasma from the same individual^120^. This approach allowed us to define an adiposity EV signature (CD63+/APMAP+/PPARG+/ADIPOQ^low^) in “primary source” ATEVs and aEVs, which reflects their adipocyte origin and functional relevance to lipid metabolism. Importantly, this adiposity EV signature was validated in bEVs (paired plasma EVs) and neat plasma from the same biological replicates. While bEVs exhibited greater variability in the expression of CD63, APMAP, PPARG, and ADIPOQ, the neat plasma largely preserved the adiposity EV signature (CD63+/APMAP+/PPARG+/ADIPOQ^low^). This consistency suggests that neat plasma EVs can serve as a reliable surrogate for tissue-specific signatures, reinforcing their utility for minimally invasive liquid biopsy applications. The integration of TIRF microscopy and SP-IRIS demonstrates the potential of our pipeline to detect and quantify low-abundance tissue-specific EVs with high precision. By bridging bulk proteomics with single-EV analysis, we established a robust framework for ADEV characterization, enabling accurate identification and validation of adipose-enriched markers.

### Challenges and future directions

Our methodology provides a foundation for developing adipose-focused EV liquid biopsies, offering a non-invasive tool for diagnosing and monitoring obesity-related diseases. By defining an adiposity EV signature (CD63+/APMAP+/PPARG+/ADIPOQ^low^) and validating it across biospecimens, we set the stage for broader applications in EV-based biomarker discovery and clinical diagnostics. This adiposity EV signature, retained even in unprocessed plasma samples, highlights its translational potential for minimally invasive liquid biopsy applications.

The distinct molecular profiles of ATEVs and aEVs uncovered in this study further suggest potential therapeutic opportunities. ATEVs, for example, could serve as therapeutic targets to modulate systemic inflammation, whereas enhancing aEV function may hold promise for addressing oxidative stress and protein processing abnormalities. Despite their potential, tissue-derived EVs represent a mere 0.2% of the total plasma EV pool, highlighting the formidable technical challenges of investigating low-abundance, tissue-derived, and cell-specific EVs. Despite these signal-to-noise limitations, adipose tissue emerges as the prominent source, contributing over 82% of tissue-derived EVs in human plasma^121^. Leveraging our next-generation, ultrasensitive single EV methodologies, we aim to define a distinctive adiposity EV signature, positioning it for potential liquid biopsy applications. In this pioneering study, we are the first to establish a comprehensive multiparametric pipeline to characterize bulk and single ADEVs from matched human visceral adipose tissue, autologous mature adipocytes, and plasma from the same bariatric patient. Notably, this research marks a breakthrough in directly detecting an adipose-specific EV signature from unprocessed plasma samples of bariatric patients. Our findings not only lay the groundwork for an obesity-focused EV liquid biopsy but also expand the therapeutic potential of novel human adipose-enriched EV as promising candidates for future translational application.

## Methods

Our study adheres to all applicable ethical regulations. The OSU Institutional Review Board (IRB) approved the study, and all participants provided written informed consent. While the study involved human subjects, they were not prospectively assigned to an intervention to evaluate the effect on health-related biological outcomes or compare diets or treatments. As a result, the study is exempt from registration in Clinical Trials.gov.

### Study participants

VAT biopsies and plasma samples were obtained from patients with 11, average age: 41.6 years old SD: 12.98, BMI: 40.97, SD: 6.00) who underwent elective bariatric surgery at The Ohio State University (OSU) Center for Minimally Invasive Surgery. Clinical data is summarized in Supplementary **Table 1**. Subjects received no compensation for participation.

### Adipose tissue dissociation and mature adipocyte culture for EV isolation

Adipose tissue samples were obtained during bariatric surgery and dissociated within an hour from procurement using collagenase digestion. The resulting cell suspension was centrifuged at 500 x g for 10 minutes to separate the mature adipocytes, the stromal vascular fraction (SVF), and the dissociation solution. The mature adipocytes were cultured for 24 hr in Dulbecco’s Modified Eagle Medium/Ham’s Nutrient Mixture F-12 (DMEM/F12), supplemented with 40% M199 basal medium with Earle’s salts and L-glutamine (MCDB201 Media), 2% Fetal Bovine Serum (FBS), 1x Penicillin/Streptomycin, 1 nanomolar (nM) dexamethasone, L-ascorbic acid 2-phosphate (LAAP) 0.1 millimolar (mM), Insulin-Transferrin-Selenium (ITS) MIX 1x, Linoleic acid-Albumin 1x, insulin 5 micrograms per microliter (µg/uL), and Triiodothyronine (T3) 1 nM.

### ADEV isolation via tangential flow filtration (TFF)

Adipose tissue dissociation media and mature adipocyte culture media were first filtered through 0.2 µm filters. Media was subsequently concentrated into 5 ml and diafiltrated using TFF with a 500 kDa filter for purification at a flow rate of 35 ml/min as previously described^122^. After TFF, the retentates were concentrated using centrifugal units (30 kDa MWCO, MilliporeSigma Amicon Ultra Centrifugal Filter Unit, Fisher Scientific) at 4000 × g for 30 min to a final volume of 100 µL.

### Plasma EV isolation via size exclusion chromatography (SEC)

Whole blood was collected in (ethylenediaminetetraacetic acid) EDTA tubes and centrifuged at 1500 x g, 10 min at RT to separate the plasma from blood cells. The plasma was then transferred to a new tube and subjected to two sequential centrifugations at 2,500 x g for 15 min at RT. The resulting platelet-depleted plasma was collected and subjected to EV isolation via SEC using the qEV single EV columns. A total of 8 fractions were collected, and fractions 1-4 were used for EV characterization. Briefly, 150 μL of platelet-depleted plasma was loaded onto the top of the qEV column, and EVs were separated based on their size and surface properties. The column was eluted with phosphate-buffered saline, and 200 μL pooled and concentrated using centrifugal filters units (30 kDa MWCO, MilliporeSigma Amicon Ultra Centrifugal Filter Unit, Fisher Scientific) at 4000 × g for 30 min to a final volume of 100 µL.

### Microfluidic resistive pulse sensing (MRPS)

Particle number concentrations and size distributions were measured by microfluidic resistive pulse sensing using the Spectradyne nCS1 instrument (Spectradyne, Torrence, CA) equipped with C-400 polydimethylsiloxane cartridges to cover a size range of approximately 65-400nm in vesicle diameters. Data analysis was performed with the nCS1 Data Analyzer (Spectradyne, Torrence, CA).

### Western blotting

EV samples were lysed in radioimmunoprecipitation assay (RIPA) buffer (Thermo Scientific) with the addition of Thermo Scientific Pierce Protease and Phosphatase Inhibitor (Thermo Scientific) for 15 min on ice. Protein concentrations were quantified using a Micro BCA™ Protein Assay Kit (Thermo Scientific). Equivalent amounts of sample proteins in Laemmli buffer (With reducing agent, 2-Mercaptoethanol (Sigma-Aldrich) were electrophoresed on 4-20% Mini-PROTEAN® TGX Stain-Free gels (Bio-rad) and then transferred onto a polyvinylidene fluoride (PVDF) membrane (Biorad). Membranes were blocked and then probed with primary antibody diluted in TBS-T overnight at 4°C and then with horseradish peroxidase conjugated secondary antibody for 1 hr at RT. Immunoreactivity was determined using enhanced chemiluminescence (ECL) solutions (Biorad) and visualized using a Biorad ChemiDoc™ MP imaging system.

### Single-particle interferometric reflectance imaging sensing (SP-IRIS)

Unchained Labs (Boston, MA, USA) silicone chips coated with tetraspanin-specific capture spots (CD63, CD9, and CD81) and a negative control spot (mouse IgG isotype). Chips were incubated for 1 hr at RT with 1×10E8 ADEVs and 1×10E7 plasma EVs diluted in a final volume of 60 µL of incubation buffer A at room temperature. After the incubation, chips were washed 3 times for 3 min on an orbital plate shaker with wash solution B. The chips were scanned with the SP-IRIS™ R200 reader (Unchained labs) by the ExoScanner software (Unchained labs, Boston, MA, USA). The data was analyzed using SP-IRISer software (Unchained labs, Boston, MA, USA).

### Transmission electron microscopy

Two 20 µL droplets of deionized (DI) water and two droplets of UranyLess EM contrast stain (Electron Microscopy Science) were placed on a parafilm surface. Prior to that, TEM grids underwent a 1-minute plasma treatment. Subsequently, 10 µL of ATEVs, aEVs, and bEVs were deposited onto the plasma-treated surface through drop-casting. The samples were allowed to incubate on the TEM surface for 1 min, followed by gentle blotting with filter paper to remove excess liquid. The TEM grids were promptly washed by dipping them into a DI water droplet, followed by blotting with filter paper. This washing step was repeated with another DI water droplet. The same technique was then repeated for the contrast stain, with each incubation lasting 22 sec. The TEM grids were carefully stored in a grid box overnight to ensure complete drying before imaging. TEM imaging was performed using a Tecnai TF-20 microscope operating at 200 kV.

### Single EV analysis using Total internal reflection fluorescence microscopy

#### Antibody functionalization of the biochip

In accordance with the methodology delineated in the preceding publication, a biochip featuring a gold-coated substrate was painstakingly prepared and subsequently functionalized using biotin-PEG-SH^92^. The functionalization of the biochip with antibodies entailed a series of specific steps. A uniform working volume of 20 μL was employed in each well to introduce various solutions. To ensure an even distribution of solutions within the wells, all incubation stages were conducted on a shaker. Before the antibody functionalization, a thorough washing of each well was conducted by pipetting DI water ten times. Following this, a solution of NeutrAvidin (NA; Thermo Fisher Scientific) at a concentration of 50 μg/mL, diluted in phosphate-buffered saline (PBS; Thermo Fisher Scientific), was added to each well. This mixture was allowed to incubate at room temperature for 1 hour to bind with the biotin motifs on the gold surface. Excess NeutrAvidin was subsequently removed by pipetting PBS ten times, and this rinsing process was repeated three times.

#### Introduction of biotinylated capture antibodies

For the introduction of biotinylated capture antibodies, the antibodies were first biotinylated using the EZ-Link micro Sulfo-NHS-biotinylation kit (Thermo Fisher Scientific). Each antibody was then diluted to a concentration of 10 μg/mL in a 1% (w/v) solution of bovine serum albumin (BSA; Sigma-Aldrich) in PBS. The prepared biotinylated capture antibody mixture was added to each well and allowed to incubate at room temperature for 1 hour. To remove excess protein, the wells were washed by pipetting PBS up and down ten times, with this rinsing process being repeated three times.

#### Molecular beacons hybridization

The molecular beacons (MBs) were diluted down to a concentration of 5 μM for each individual MB, using 12.5 × Tris EDTA (TE) buffer (Sigma-Aldrich), which was diluted in deionized (DI) water. This dilution served the dual purpose of stabilizing the MBs and permeabilizing the RNA-encasing membrane. The resulting mixture of MBs was subsequently diluted by a factor of 25 within the biofluid sample, and the biochip was then subjected to a 2-hr incubation period at 37°C, creating an environment devoid of light to facilitate the hybridization of the MBs with the target RNA.

#### Capture of EVs on biochip surface

In each well, a solution of BSA in PBS, with a concentration of 3% (w/v), underwent a 1-hour incubation at room temperature. This step aimed to prevent nonspecific EV capture and maintain single-vesicle resolution. Following the removal of BSA, the biofluid samples were then introduced into the wells and allowed to incubate for 2 hours at room temperature under dark conditions. PBS was introduced into the wells for a 5-minute incubation period, followed by pipetting PBS up and down ten times to eliminate excessive EVs and unhybridized MBs. This rinsing process was repeated four times within a dimly lit environment.

#### Detection antibodies targeting membrane proteins

Initially, a solution of BSA at a concentration of 3% (w/v) in PBS was added to each well and incubated at room temperature for one hour. This step aimed to prevent the nonspecific binding of the fluorophore-conjugated antibodies (detection antibodies). After the BSA solution was removed, a solution containing detection antibodies at a concentration of 1 µg/mL was prepared in 10% (w/v) normal goat serum (NGS; Thermo Fisher Scientific). Subsequently, the wells were subjected to a one-hour incubation at room temperature in a dark setting. To eliminate excess detection antibodies, PBS was added to the wells and allowed to incubate for 5 minutes. Following this, gentle pipetting up and down of the solution was carried out ten times. The rinsing procedure was repeated thrice within a dimly lit environment.

#### EV proteomics digestion

EVs were digested using Protifi S-trap^TM^ and the published protocol was adapted to fit the project needs^123^. Briefly, samples were digested in lysis buffer consisting of 10% Sodium Dodecyl Sulfate (SDS) with 100 millimolar (mM) tetraethylammonium bromide (TEAB). Samples were sonicated and then clarified by centrifugation. Protein concentration was determined using NanoDrop A280 (Thermo Fisher Scientific). Disulfide bonds were reduced by addition of dithiothreitol (DTT), final concentration 20 mM, and incubation at 95LC for 15 min. Free cysteines were alkylated by the addition of iodoacetamide at a final concentration of 40 mM, followed by incubation in the dark for 30 minutes at room temperature (RT). Unreacted iodoacetamide was removed by centrifugation, after which the supernatant was collected. Samples were mixed with binding buffer in a 1:6 ratio of sample to buffer volume. The resulting solution was loaded onto a Protifi S-trap^TM^ micro spin column in increments of 300 μL, and centrifuged until all protein was bound to the column. 20 μL of digestion buffer was added, containing Trypsin Gold protease, mass spectrometry grade (Promega, ref V528A) in a 1:10 enzyme-to-protein ratio for each sample. Trypsin was resuspended in 50mM triethylammonium bicarbonate (TEAB) at a 1mg/mL concentration for the stock solution (Sigma ref T7408). Samples were kept at 37C overnight and eluted with sequential washes of 50 mM TEAB in water, 0.2% formic acid in water, and 0.2% formic acid in 50:50 water/acetonitrile. Samples were dried and resuspended in 0.1% formic acid in water for downstream analysis.

#### LC-MS/MS analysis

Bottom-up proteomics data were collected on a Bruker Scientific (Billerica, MA) commercial timsTOF Pro instrument coupled to a Bruker Scientific nanoElute LC. For each sample, 200 ng was loaded. Samples were run in data-dependent acquisition (DDA) mode, using parallel accumulation-serial fragmentation (PASEF). The LC column was 25 cm long with 1.7 μm C18 particles (IonOpticks), and samples were run at a rate of 0.40 μL. Funrich^124^ software was used to generate the Venn diagram. This was accomplished by taking the total lists of proteins identified in aEVs and ATEVs separately and uploading them to Funrich as protein lists. The third group is the Vesiclepedia database^125^, a list of proteins implicated in EVs which have been previously reported in the literature.

/min, using a linear reverse phase 120-minute gradient with a starting concentration of 2.0% acetonitrile (ACN) with 0.1% formic acid (FA) and ending concentration of 80.0% ACN with 0.1% FA over a period of 110 minutes, with a 10-minute ending hold. The equilibration time was estimated at 3.60 minutes. For MS/MS analysis, the mass range was set as 100 to 1700 m/z, with 10 MS/MS scans, dynamic exclusion time of 0.4 min, and collision-induced dissociation (CID) energy starting at 20.00 eV and ending at 59.00 eV.

#### Proteomics data processing and analysis

TimsTOF PASEF datasets were processed using FragPipe (v. 19.1) in conjunction with the MS/MS search engine MSFragger (v. 3.7) and IonQuant (v. 1.8.10). The Uniport human reviewed database was downloaded and used as the input fasta file for spectral mapping. Common contaminants and decoys were added using FragPipe software. The fraction of decoys was set to 50% and was generated by reversing sequences from the target database. This target-decoy search was used to control the false discovery rate (FDR)^126^. The precursor mass tolerance was 20 ppm, and the enzyme was set to trypsin with 2 missed cleavages. Peptide length was set to 7– 50, and peptide mass range was 500 – 5,000. Variable modifications were set to include methionine oxidation (+15.9949) and N-terminal acetylation (+42.0106). Carbamidomethylation on cysteine residues from iodoacetamide treatment (+57.021464) was set as a fixed modification. The number of variable modifications on a given peptide was set to 3. Percolator was used for peptide-spectrum match (PSM) validation, and MS1 quantification was run using match between runs (MBR) with an FDR of 0.01. Intensity was normalized across runs, using a log-ratio approach where the run with the highest number of ions is used as a reference. Peptide-protein uniqueness was set to unique + razor, where unique peptide refers to the number of PSMs that only map to the given protein, and razor refers to the total number of PSMs that in support of a given protein.

FragPipe results were exported as TSV (tab separated value) files, compatible with Excel for downstream processing. For identified proteins, precursor/MS1 intensity values were used as quantitative markers of protein expression. Intensity is defined by IonQuant software as the summed volume of the 0, +1 and +2 isotopes, where the volume is from ion mobility and retention time information. Missing values were replaced with an appropriate low number (5000). For individual protein identifications, values that were > 50% missing from either ATEV or aEV groups were removed. Values that were not consistent (i.e., sometimes present, sometimes missing) across a single group (ATEV or aEV) were also removed. The data were log base 10 transformed and quantile normalized. A two-sided t test was used to compare ATEVs to aEVs. P-values were FDR-corrected using the Benjamini-Hochberg method. A one-sided t-test was used to look at the ratios of donor-matched aEVs to ATEVs. FDR of 0.05 was considered significant.

Proteome FragPipe outputs were further processed using the DAVID (Database for Annotation, Visualization, and Integrated Discovery) bioinformatics tool^127,128^.The GO (Gene Ontology) pathway and KEGG (Kyoto Encyclopedia of Genes and Genomes) pathway analyses were conducted. The DAVID tool allowed us to select the statistical analysis method, and we opted to use the FDR as a threshold for identifying significant pathway enrichments. . Funrich^124^ software was used to generate the Venn diagram. This was accomplished by taking the total lists of proteins identified in aEVs and ATEVs separately and uploading them to Funrich as protein lists. The third group is the Vesiclepedia database^125^, a list of proteins implicated in EVs which have been previously reported in the literature.

### Protein-Protein interaction analysis

To investigate the molecular interactions among the differentially expressed proteins identified in this study, we utilized the STRING database (version 12.0)^129^ and Cytoscape (version 3.10.2.)^130,131^ . Proteins were input into STRING using *Homo sapiens* as the reference organism, with interaction sources including experimental data, curated databases, text mining, and co-expression. A high-confidence interaction threshold (minimum score 0.7) was applied to generate a robust protein-protein interaction (PPI) network, encompassing both direct and indirect interactions. The resulting network was imported into Cytoscape^130,131^ via the STRING application^129^ for visualization and further analysis.

Using Cytoscape application, network topology metrics such as degree centrality and betweenness centrality were calculated to identify hub proteins^131^. Functional enrichment analysis of Gene Ontology (GO) terms and KEGG pathways was performed using STRING-integrated tools^129^. Cluster analysis using the MCODE plugin^132^ identified densely connected modules, highlighting potential protein complexes and key functional groups. Additionally, the network was refined by integrating literature-based interactions to enhance biological relevance.

## Data availability

All data generated or analyzed during this study are included in this manuscript and its supplementary information file. Further inquiries can be directed to the corresponding author.

## Author contributions

M.D.H., P.L.P., W.H., E.R., S.M.M, conceptualization.

E.R., S.M.M. study design, and funding acquisition.

M.D.H., J.G., P.L.P., D.S., K.T.N., B.L.B., J.K. S.N., S.A.B., B.J.N., V.W., W.H. investigation and performed research.

M.D.H., J.G., P.L.P., K.T.N., J.K. organization and data analysis.

M.D.H., P.L.P., S.M.M completed the original draft of the manuscript.

M.D.H., E.R., S.M.M. data interpretation, revision, and manuscript correction. All authors approved the final version of the manuscript.

## Additional information

Correspondence and requests for materials should be addressed to Senior Corresponding author.

## Acknowledgments

We would like to thank William Ray for his illustrative media expertise in designing the figures for this project. Drs. Kristy Townsend and Kristin Stanford provided technical input for adipose tissue processing. SMM was supported by intramural funding from the Abigail Wexner Research Institute at Nationwide Children’s Hospital and NINDS (5K12NS098482-05). ER was supported by National Institutes of Health (NIH) grants UG3/UH3TR002884 and U18TR003807. JG was supported by an institutional pilot grant from the Metabolic and Diabetes Research Center. The content is solely the responsibility of the authors and does not necessarily represent the official views of the National Institutes of Health.

