## Supplementary material for "Novel multiparametric bulk and single extracellular vesicle pipeline for adipose cell-specific biomarker discovery in paired human biospecimens": Suppl Material

**Running Title:** Human adipose extracellular vesicles

***Corresponding Author:** Setty Magaña, MD, PhD

Address:	Nationwide Children’s Hospital

Department of Pediatrics, Division of Neurology

700 Children’s Drive

Columbus, OH 43205

**Supplementary Figure 1**. Tetraspanin colocalization analysis for ATEV (left) and aEV (right). Fluorescently-labelled anti-CD63, anti-CD81, anti-CD9 were used for particle detection after particle capture with CD63, CD81, and CD9. Capture antibody is indicated in the upper right corner of each graph. ATEVs showed similar CD9 and CD81 co-expression levels, while aEVs CD9 positive particles demonstrated less degree of co-expression with CD81.
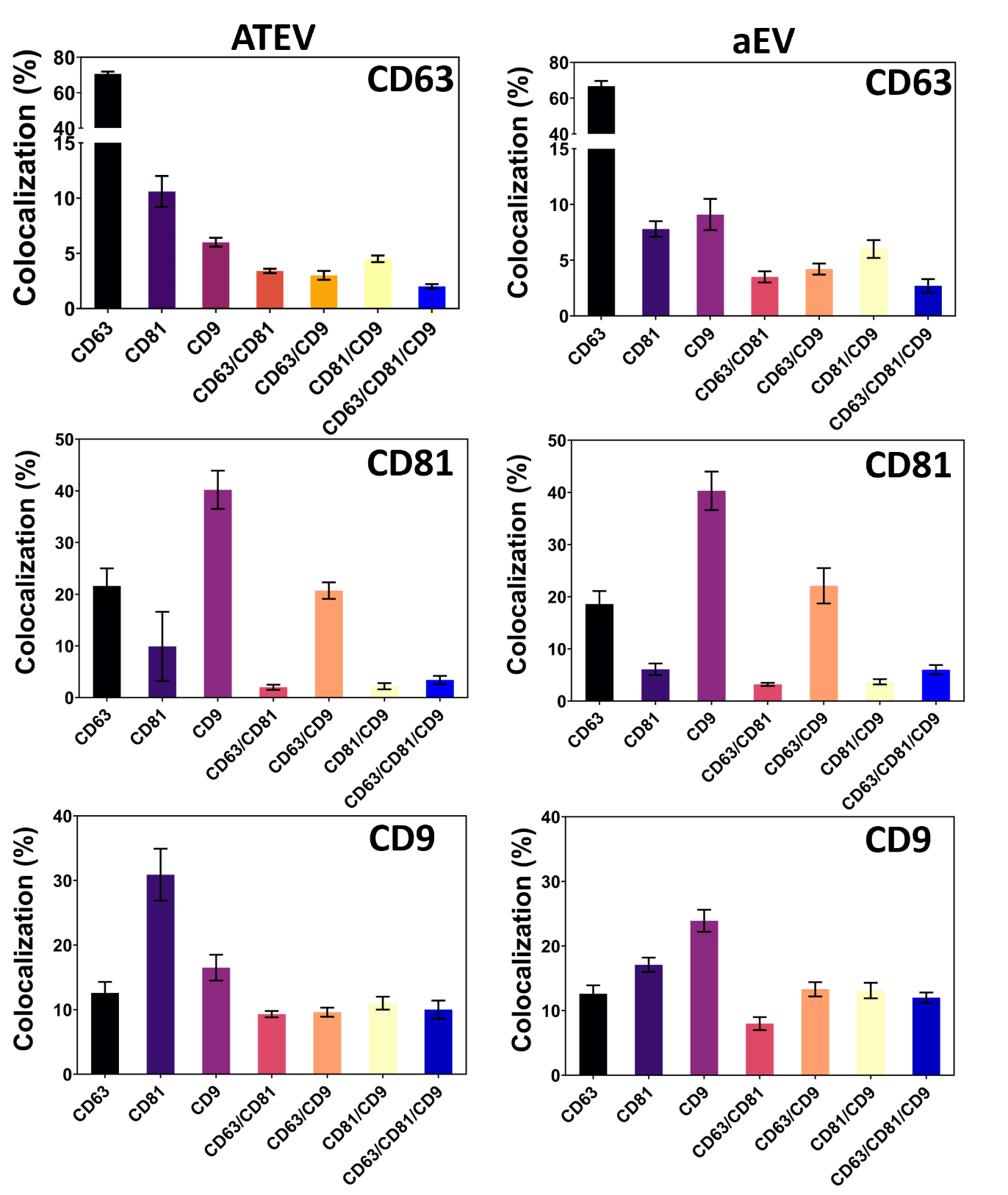

**Supplementary Figure 2**. Tetraspanin colocalization analysis for plasma EVs (bEVs). Fluorescently-labelled anti-CD63, anti-CD81, anti-CD9 were used for particle detection after particle capture with CD63, CD81, and CD9. Capture antibody is indicated in the upper right corner of each graph. Colocalization analysis demonstrated a high percentage of CD63+/CD81+ double-positive bEVs.

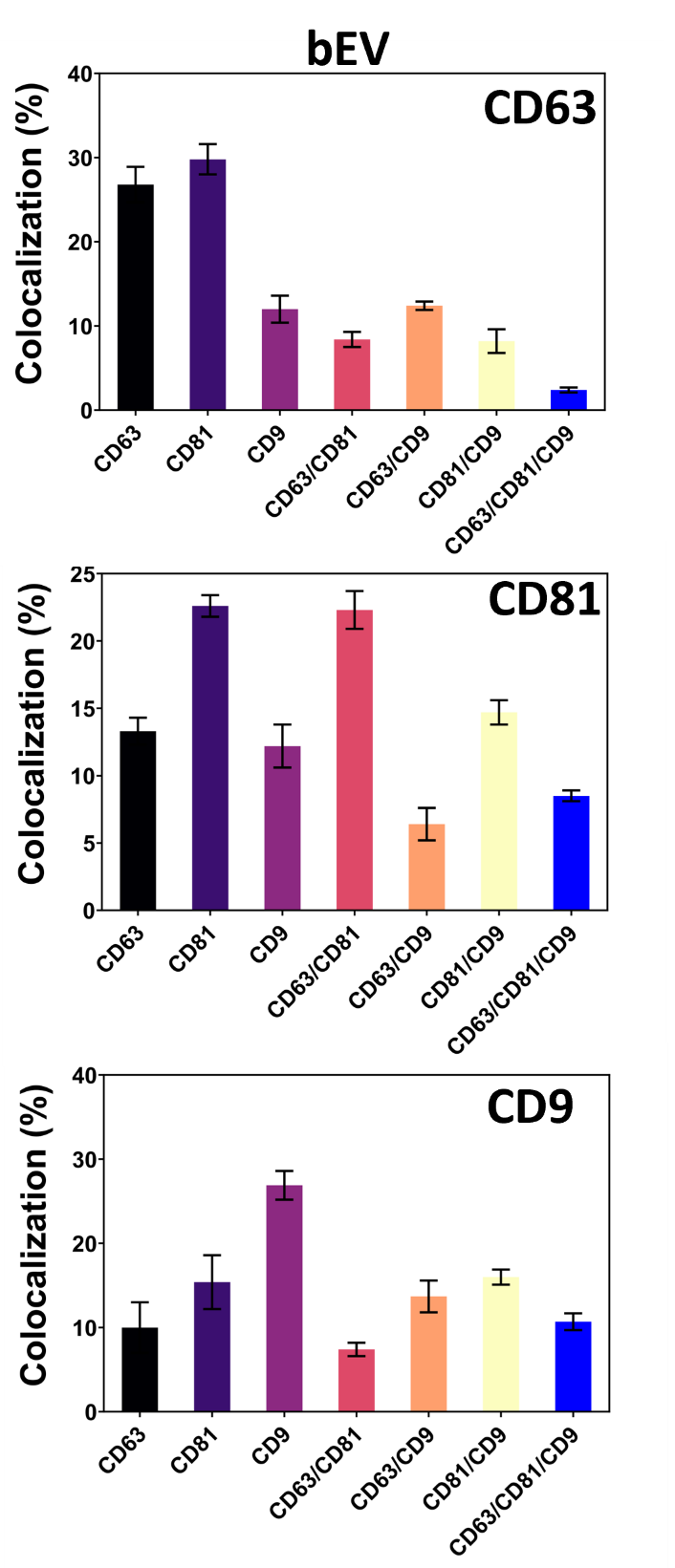

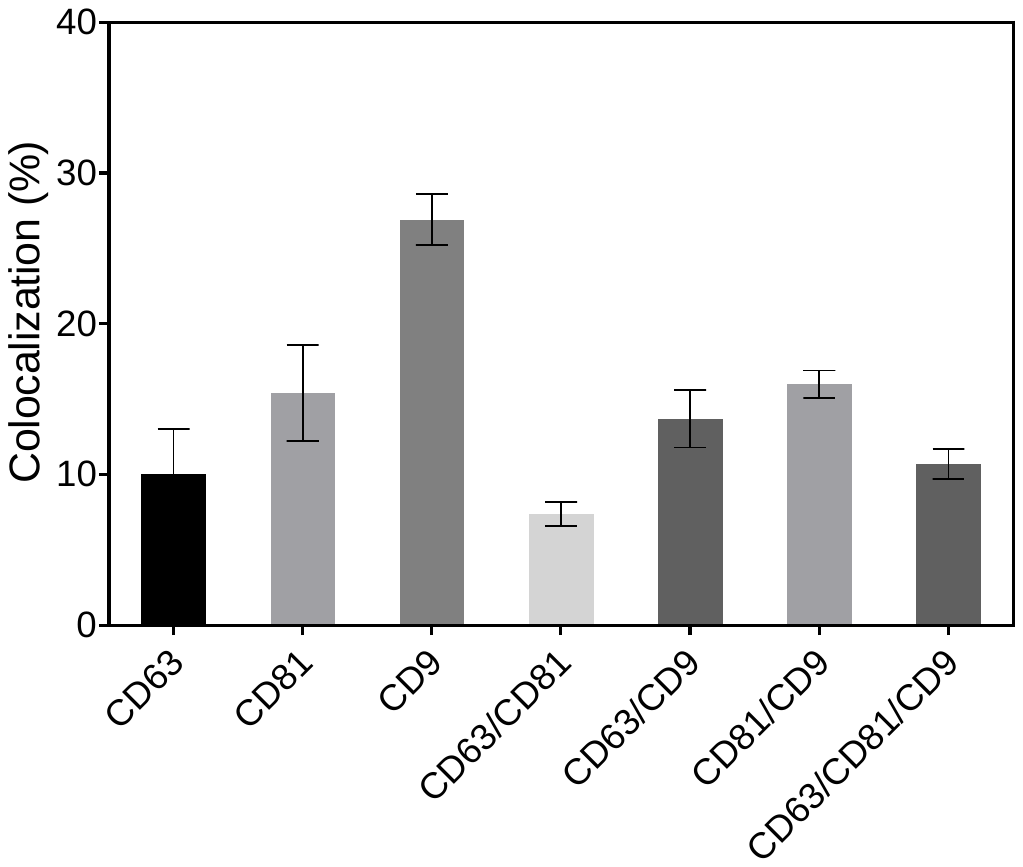

**Supplementary Figure 3**. **An interaction map for LAMC1 and its functional cellular partners.** (a) protein-protein interaction map (b) Biological process (GO) enrichment (c) Molecular function (GO) enrichment (d) Cellular function (GO) enrichment (e) KEGG pathways enrichment. Protein interaction networks were generated using STRING database integrated with Cytoscape software (v3.10.2) with a minimum interaction score threshold of 0.4 for significantly differentially expressed proteins. Edges indicate protein-protein interactions, represented by different colors: known Interactions (Curated Databases: Light blue line, Experimentally Determined: Pink line), Predicted Interactions (Gene Neighborhood: Green line, Gene Fusions: Red line), Gene Co-occurrence: Blue line, Others (Text mining: Yellow line, Co-expression: Black line, Protein Homology: Lavender line).

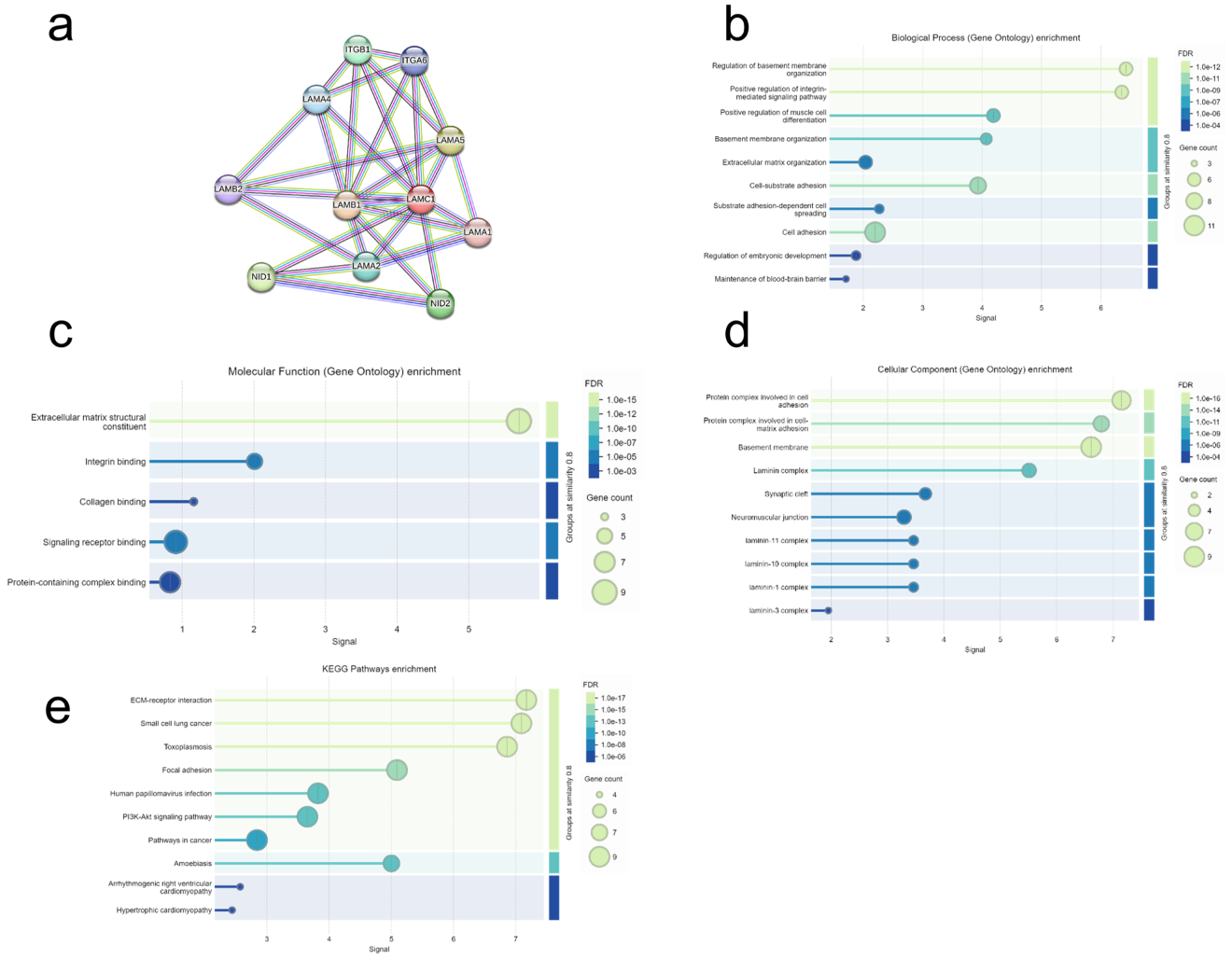

**Supplementary Figure 4. An interaction map for LAMB2 and its functional cellular partners.** (a) protein-protein interaction map (b) Biological process (GO) enrichment (c) Molecular function (GO) enrichment (d) Cellular function (GO) enrichment (e) KEGG pathways enrichment. Protein interaction networks were generated using STRING database integrated with Cytoscape software (v3.10.2) with a minimum interaction score threshold of 0.4 for significantly differentially expressed proteins. Edges indicate protein-protein interactions, represented by different colors: known Interactions (Curated Databases: Light blue line, Experimentally Determined: Pink line), Predicted Interactions (Gene Neighborhood: Green line, Gene Fusions: Red line), Gene Co-occurrence: Blue line, Others (Text mining: Yellow line, Co-expression: Black line, Protein Homology: Lavender line).

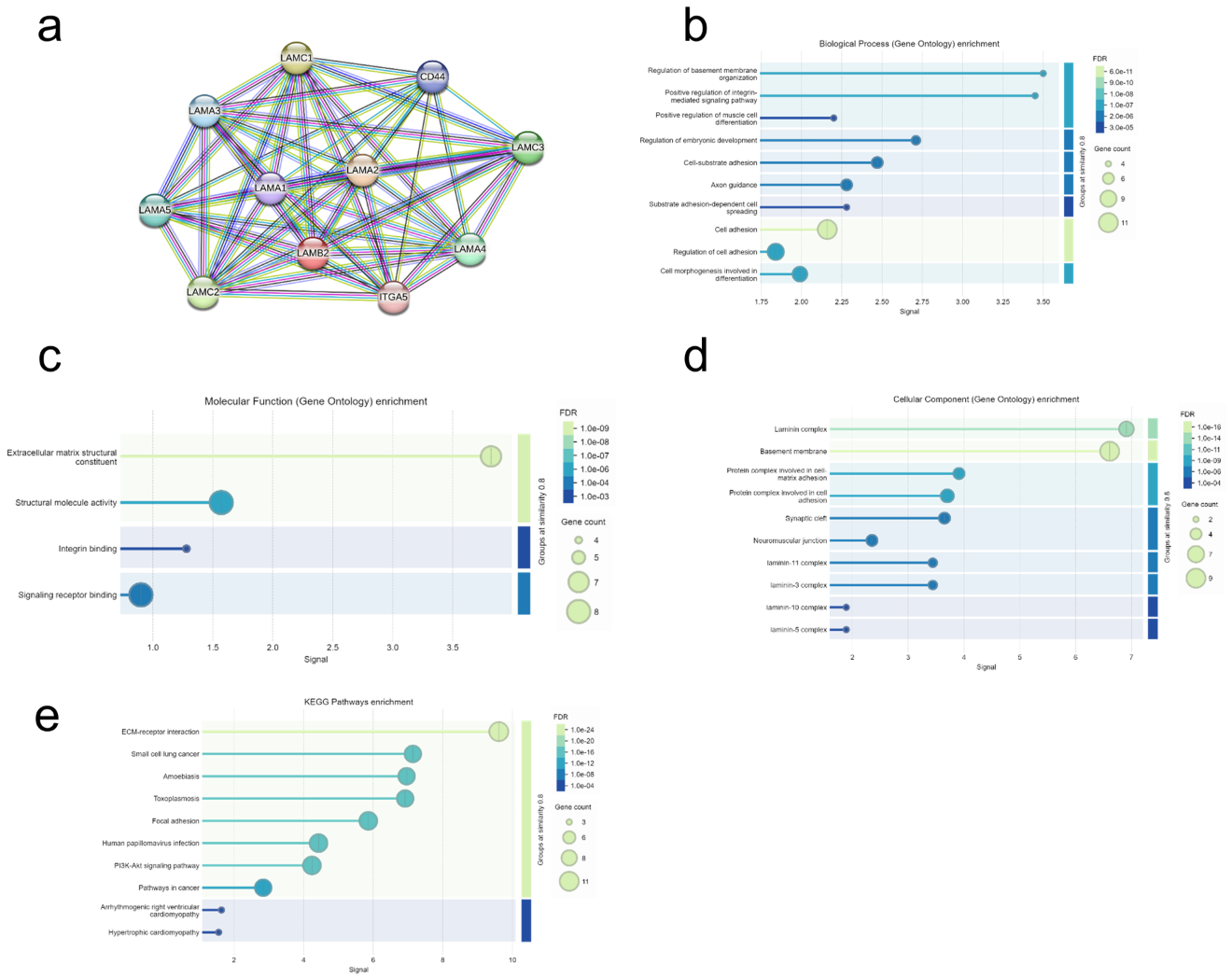

**Supplementary Figure 5. An interaction map for CYB5R3 and its functional cellular partners.** (a) protein-protein interaction map (b) Biological process (GO) enrichment (c) Molecular function (GO) enrichment (d) Cellular function (GO) enrichment (e) KEGG pathways enrichment. Protein interaction networks were generated using STRING database integrated with Cytoscape software (v3.10.2) with a minimum interaction score threshold of 0.4 for significantly differentially expressed proteins. Edges indicate protein-protein interactions, represented by different colors: known Interactions (Curated Databases: Light blue line, Experimentally Determined: Pink line), Predicted Interactions (Gene Neighborhood: Green line, Gene Fusions: Red line), Gene Co-occurrence: Blue line, Others (Text mining: Yellow line, Co-expression: Black line, Protein Homology: Lavender line).

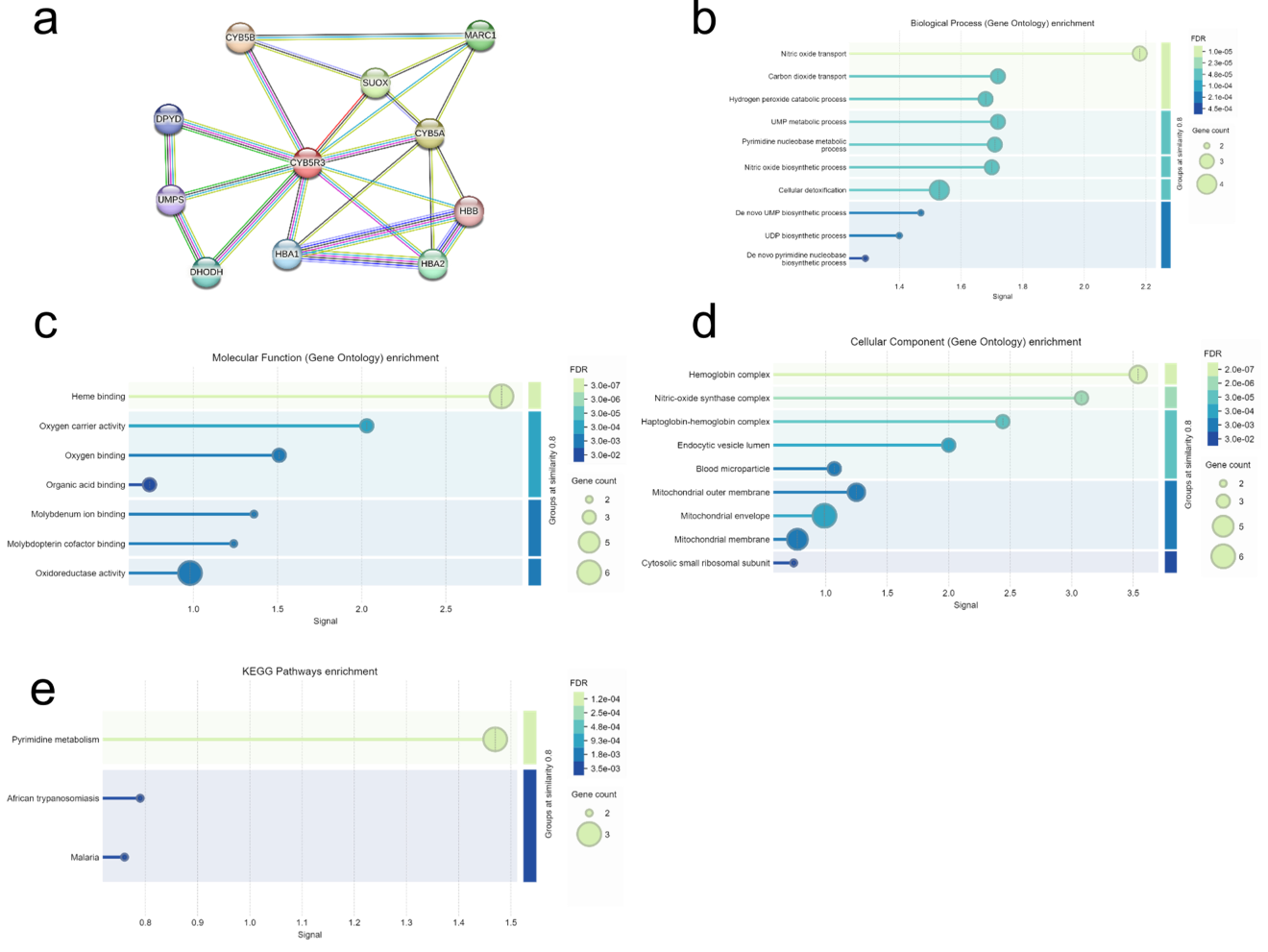

**Supplementary Figure 6. An interaction map for PRXL2A and its functional cellular partners.** (a) protein-protein interaction map (b) KEGG pathways enrichment. Protein interaction networks were generated using STRING database integrated with Cytoscape software (v3.10.2) with a minimum interaction score threshold of 0.4 for significantly differentially expressed proteins. Edges indicate protein-protein interactions, represented by different colors: known Interactions (Curated Databases: Light blue line, Experimentally Determined: Pink line), Predicted Interactions (Gene Neighborhood: Green line, Gene Fusions: Red line), Gene Co-occurrence: Blue line, Others (Text mining: Yellow line, Co-expression: Black line, Protein Homology: Lavender line).

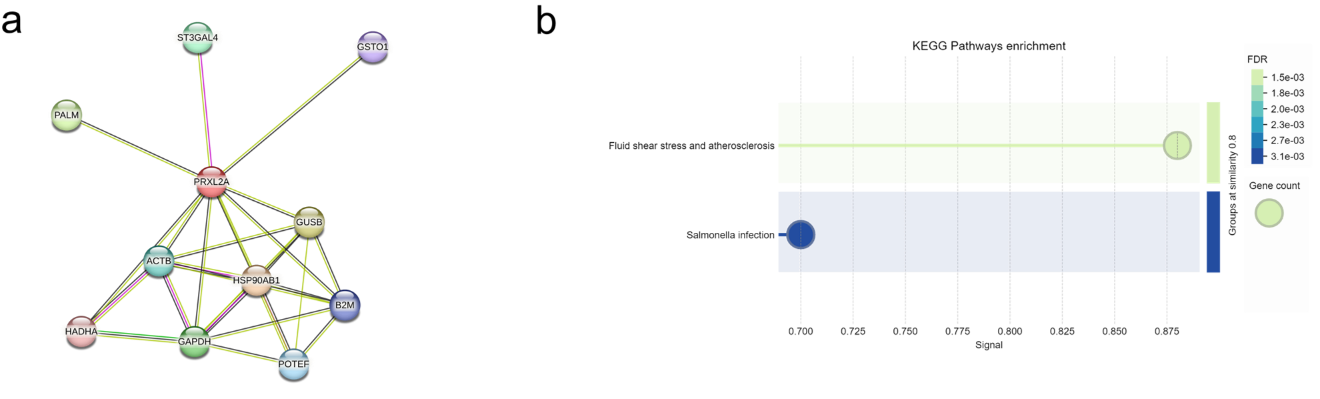

**Supplementary Figure 7. An interaction map for RTN3 and RTN4 and its functional cellular partners.** (a) protein-protein interaction map (b) Biological process (GO) enrichment (c) Molecular function (GO) enrichment (d) Cellular function (GO) enrichment (e) KEGG pathways enrichment. Protein interaction networks were generated using STRING database integrated with Cytoscape software (v3.10.2) with a minimum interaction score threshold of 0.4 for significantly differentially expressed proteins. Edges indicate protein-protein interactions, represented by different colors: known Interactions (Curated Databases: Light blue line, Experimentally Determined: Pink line), Predicted Interactions (Gene Neighborhood: Green line, Gene Fusions: Red line), Gene Co-occurrence: Blue line, Others (Text mining: Yellow line, Co-expression: Black line, Protein Homology: Lavender line).

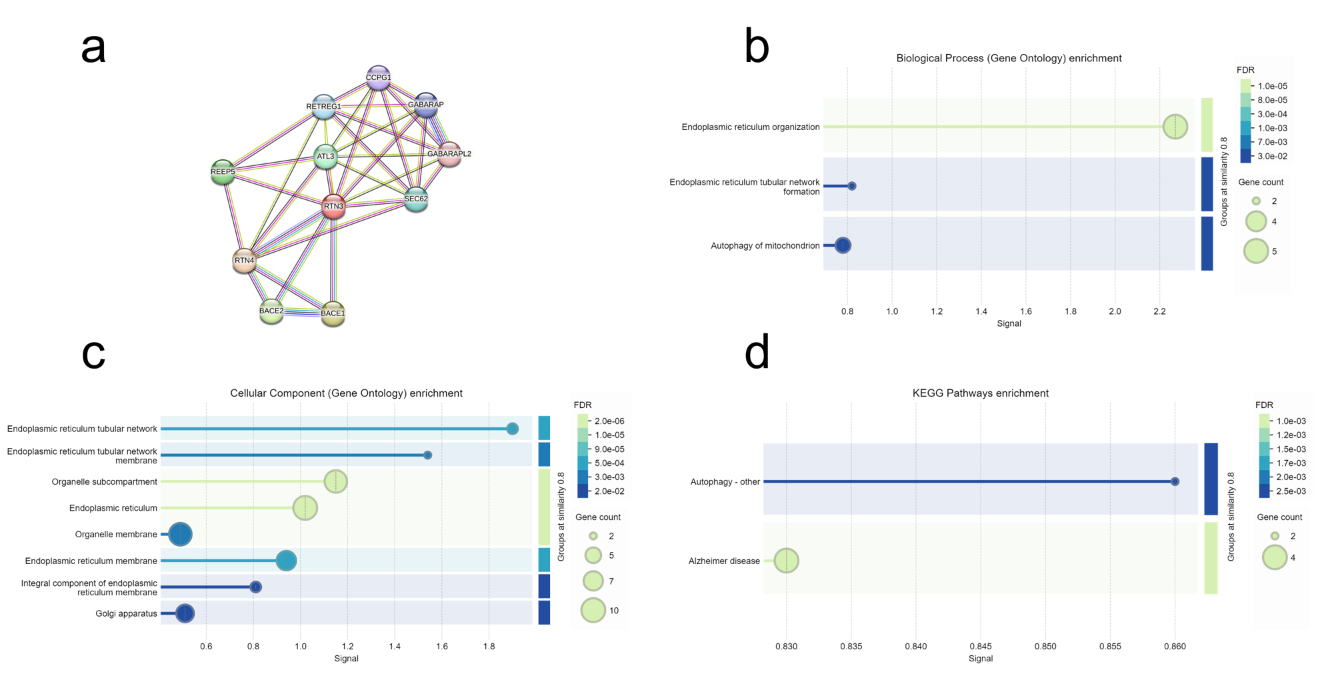

**Supplementary Figure 8. An interaction map for GNAI2 and its functional cellular partners.** (a) protein-protein interaction map (b) Biological process (GO) enrichment (c) Molecular function (GO) enrichment (d) Cellular function (GO) enrichment (e) KEGG pathways enrichment. Protein interaction networks were generated using STRING database integrated with Cytoscape software (v3.10.2) with a minimum interaction score threshold of 0.4 for significantly differentially expressed proteins. Edges indicate protein-protein interactions, represented by different colors: known Interactions (Curated Databases: Light blue line, Experimentally Determined: Pink line), Predicted Interactions (Gene Neighborhood: Green line, Gene Fusions: Red line), Gene Co-occurrence: Blue line, Others (Text mining: Yellow line, Co-expression: Black line, Protein Homology: Lavender line).

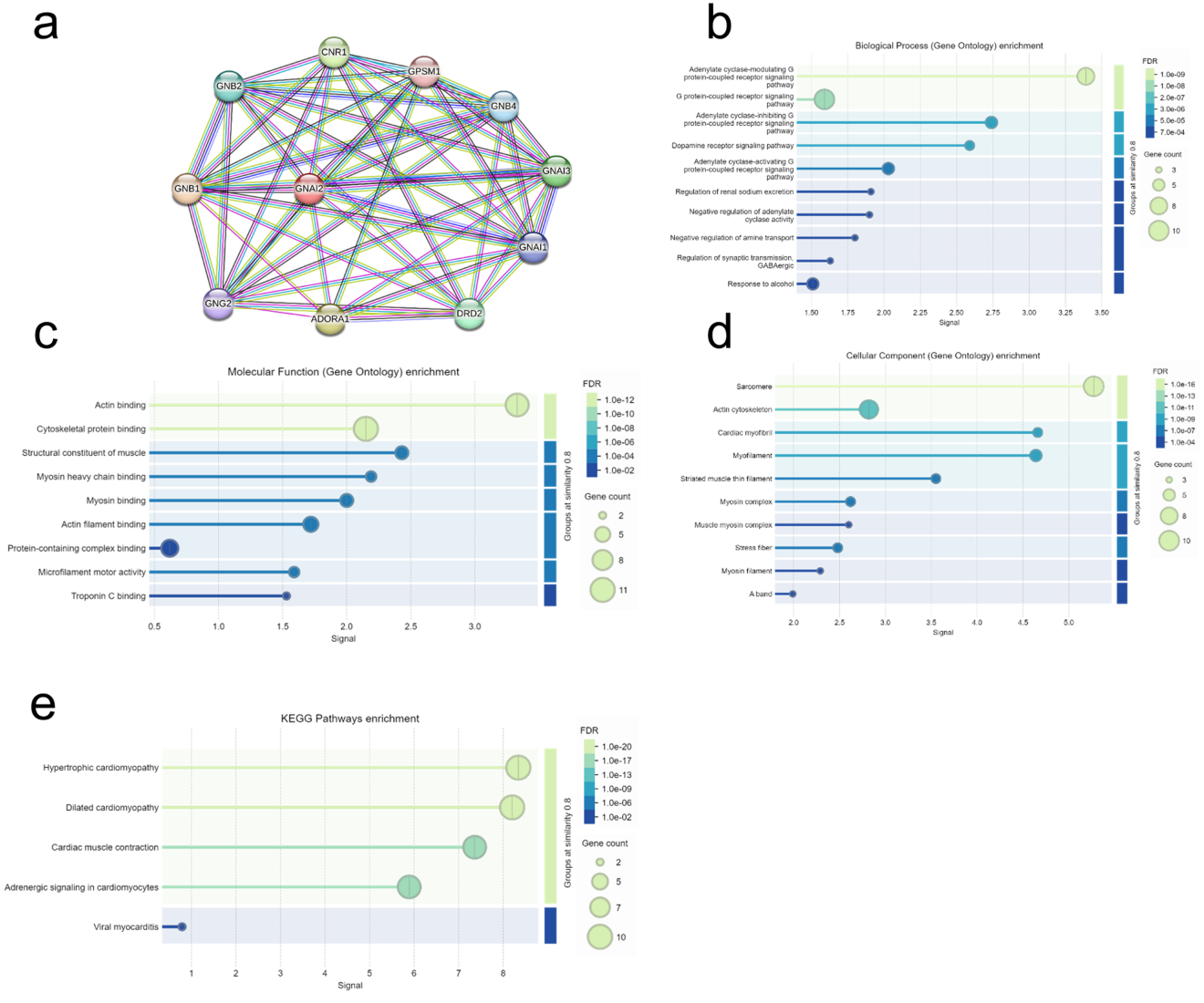

**Supplementary Figure 9. An interaction map for FTL and its functional cellular partners.** (a) protein-protein interaction map (b) Biological process (GO) enrichment (c) Molecular function (GO) enrichment (d) Cellular function (GO) enrichment (e) KEGG pathways enrichment. Protein interaction networks were generated using STRING database integrated with Cytoscape software (v3.10.2) with a minimum interaction score threshold of 0.4 for significantly differentially expressed proteins. Edges indicate protein-protein interactions, represented by different colors: known Interactions (Curated Databases: Light blue line, Experimentally Determined: Pink line), Predicted Interactions (Gene Neighborhood: Green line, Gene Fusions: Red line), Gene Co-occurrence: Blue line, Others (Text mining: Yellow line, Co-expression: Black line, Protein Homology: Lavender line).

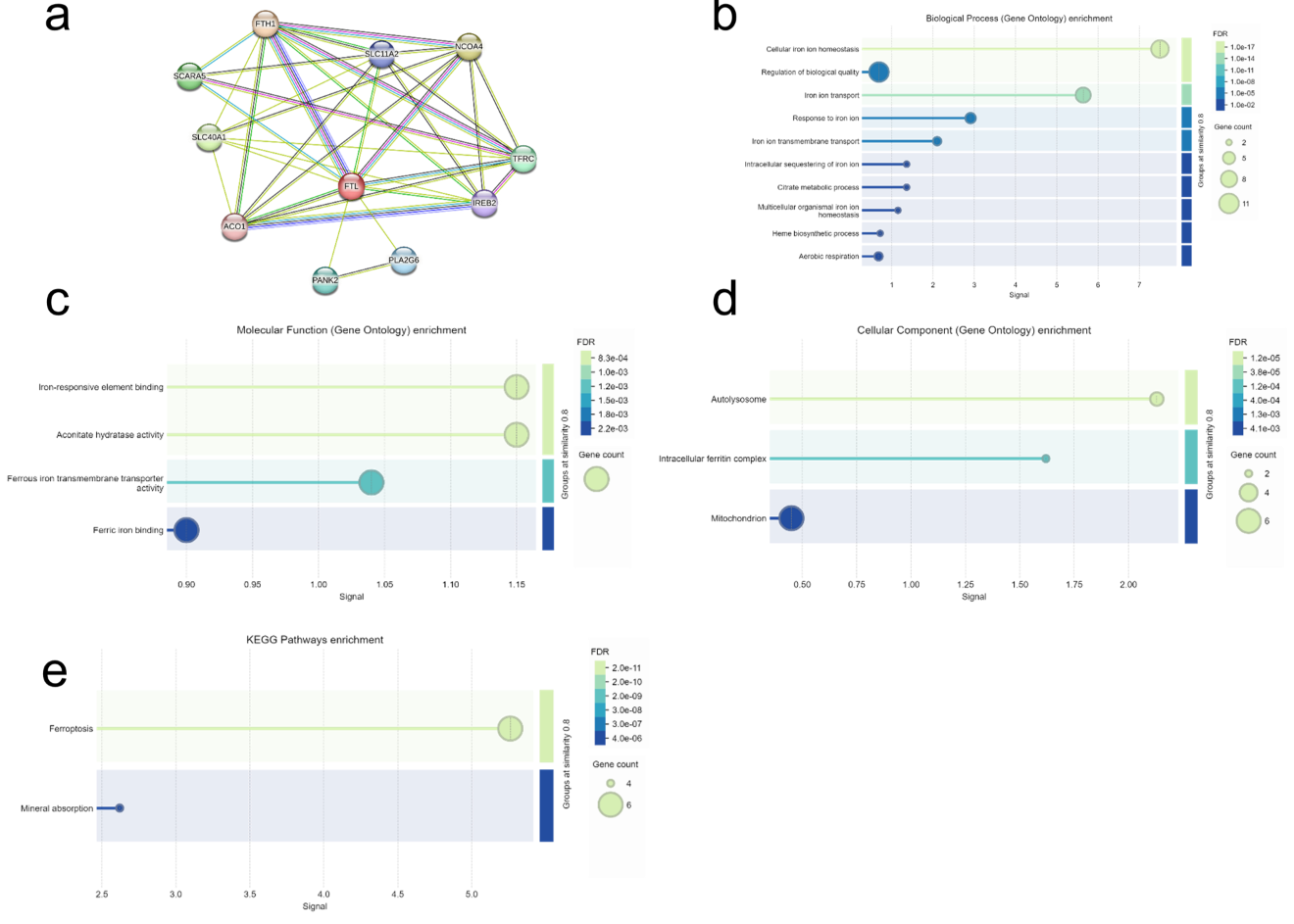

**Supplementary Figure 10. An interaction map for CLTC and its functional cellular partners.** (a) protein-protein interaction map (b) Biological process (GO) enrichment (c) Molecular function (GO) enrichment (d) Cellular function (GO) enrichment (e) KEGG pathways enrichment. Protein interaction networks were generated using STRING database integrated with Cytoscape software (v3.10.2) with a minimum interaction score threshold of 0.4 for significantly differentially expressed proteins. Edges indicate protein-protein interactions, represented by different colors: known Interactions (Curated Databases: Light blue line, Experimentally Determined: Pink line), Predicted Interactions (Gene Neighborhood: Green line, Gene Fusions: Red line), Gene Co-occurrence: Blue line, Others (Text mining: Yellow line, Co-expression: Black line, Protein Homology: Lavender line).

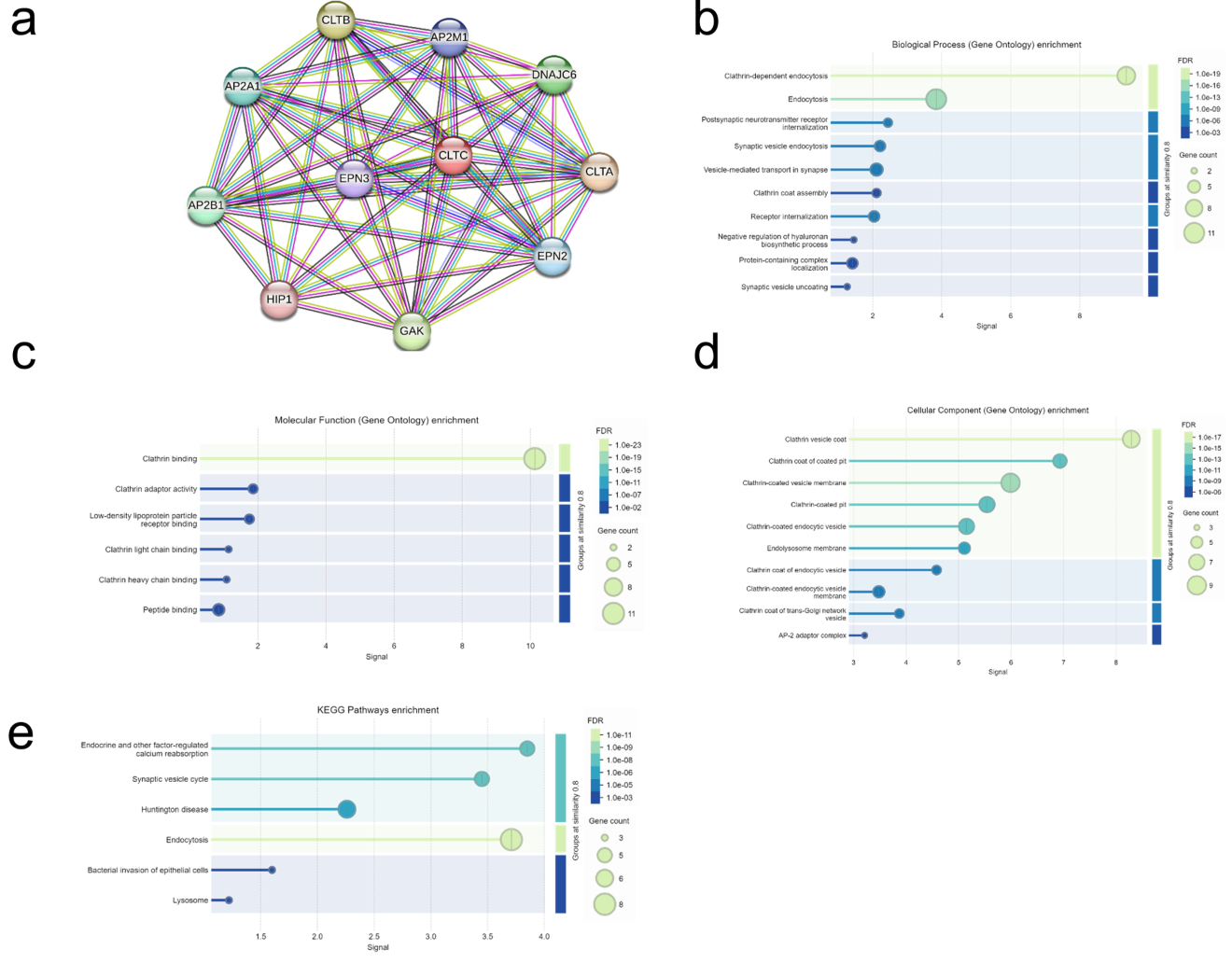

**Supplementary Figure 11. An interaction map for ATF1 and its functional cellular partners.** (a) protein-protein interaction map (b) Biological process (GO) enrichment (c) Molecular function (GO) enrichment (d) Cellular function (GO) enrichment (e) KEGG pathways enrichment. Protein interaction networks were generated using STRING database integrated with Cytoscape software (v3.10.2) with a minimum interaction score threshold of 0.4 for significantly differentially expressed proteins. Edges indicate protein-protein interactions, represented by different colors: known Interactions (Curated Databases: Light blue line, Experimentally Determined: Pink line), Predicted Interactions (Gene Neighborhood: Green line, Gene Fusions: Red line), Gene Co-occurrence: Blue line, Others (Text mining: Yellow line, Co-expression: Black line, Protein Homology: Lavender line).

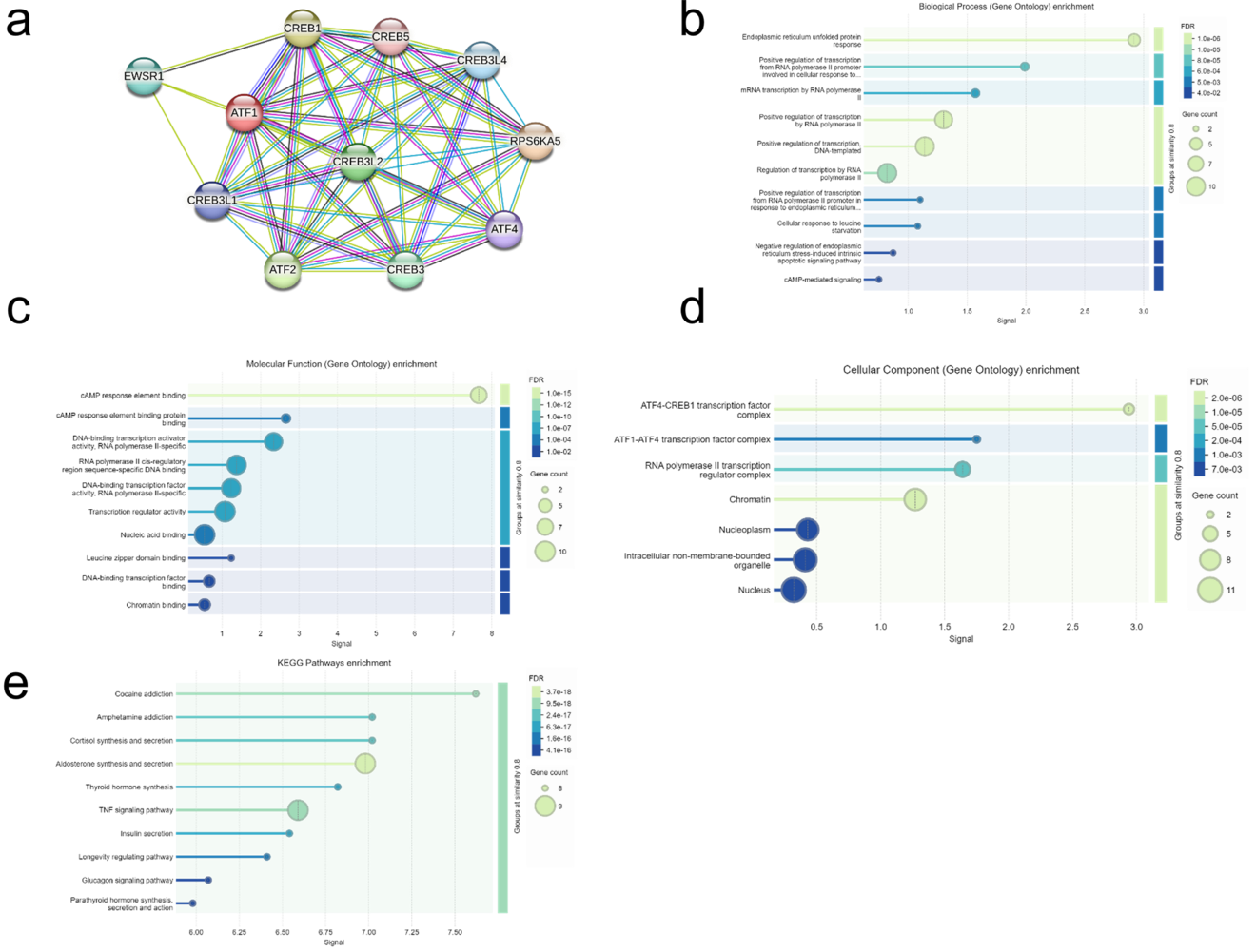

**Supplementary Figure 12. An interaction map for ACTC1 and its functional cellular partners.** (a) protein-protein interaction map (b) Biological process (GO) enrichment (c) Molecular function (GO) enrichment (d) Cellular function (GO) enrichment (e) KEGG pathways enrichment. Protein interaction networks were generated using STRING database integrated with Cytoscape software (v3.10.2) with a minimum interaction score threshold of 0.4 for significantly differentially expressed proteins. Edges indicate protein-protein interactions, represented by different colors: known Interactions (Curated Databases: Light blue line, Experimentally Determined: Pink line), Predicted Interactions (Gene Neighborhood: Green line, Gene Fusions: Red line), Gene Co-occurrence: Blue line, Others (Text mining: Yellow line, Co-expression: Black line, Protein Homology: Lavender line).

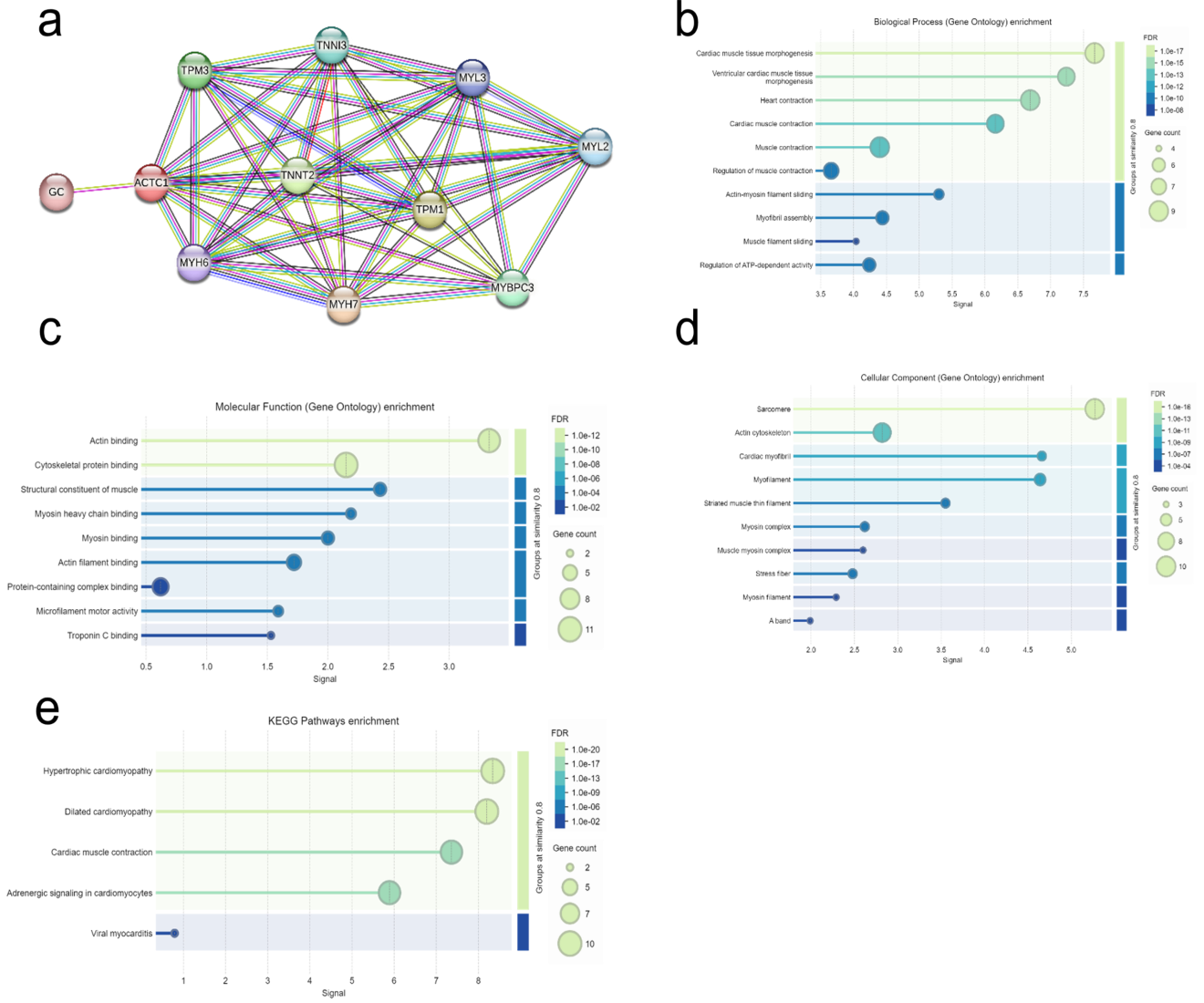

**Supplementary Figure 13. An interaction map for HSPG2 and its functional cellular partners.** (a) protein-protein interaction map (b) Biological process (GO) enrichment (c) Molecular function (GO) enrichment (d) Cellular function (GO) enrichment (e) KEGG pathways enrichment. Protein interaction networks were generated using STRING database integrated with Cytoscape software (v3.10.2) with a minimum interaction score threshold of 0.4 for significantly differentially expressed proteins. Edges indicate protein-protein interactions, represented by different colors: known Interactions (Curated Databases: Light blue line, Experimentally Determined: Pink line), Predicted Interactions (Gene Neighborhood: Green line, Gene Fusions: Red line), Gene Co-occurrence: Blue line, Others (Text mining: Yellow line, Co-expression: Black line, Protein Homology: Lavender line).

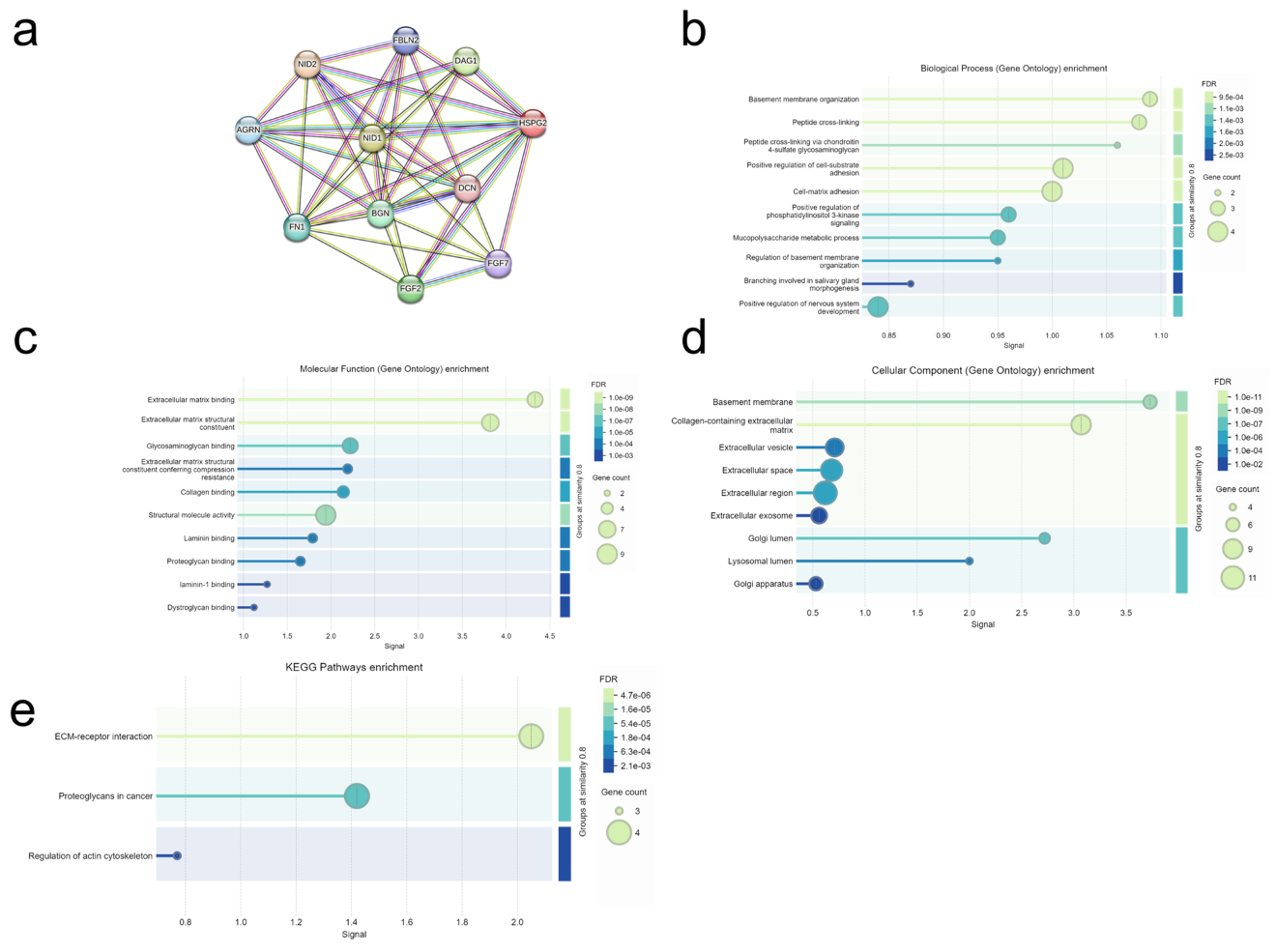

**Supplementary Figure 14. An interaction map for H2AC20 and its functional cellular partners.** (a) protein-protein interaction map (b) Biological process (GO) enrichment (c) Molecular function (GO) enrichment (d) Cellular function (GO) enrichment (e) KEGG pathways enrichment. Protein interaction networks were generated using STRING database integrated with Cytoscape software (v3.10.2) with a minimum interaction score threshold of 0.4 for significantly differentially expressed proteins. Edges indicate protein-protein interactions, represented by different colors: known Interactions (Curated Databases: Light blue line, Experimentally Determined: Pink line), Predicted Interactions (Gene Neighborhood: Green line, Gene Fusions: Red line), Gene Co-occurrence: Blue line, Others (Text mining: Yellow line, Co-expression: Black line, Protein Homology: Lavender line).

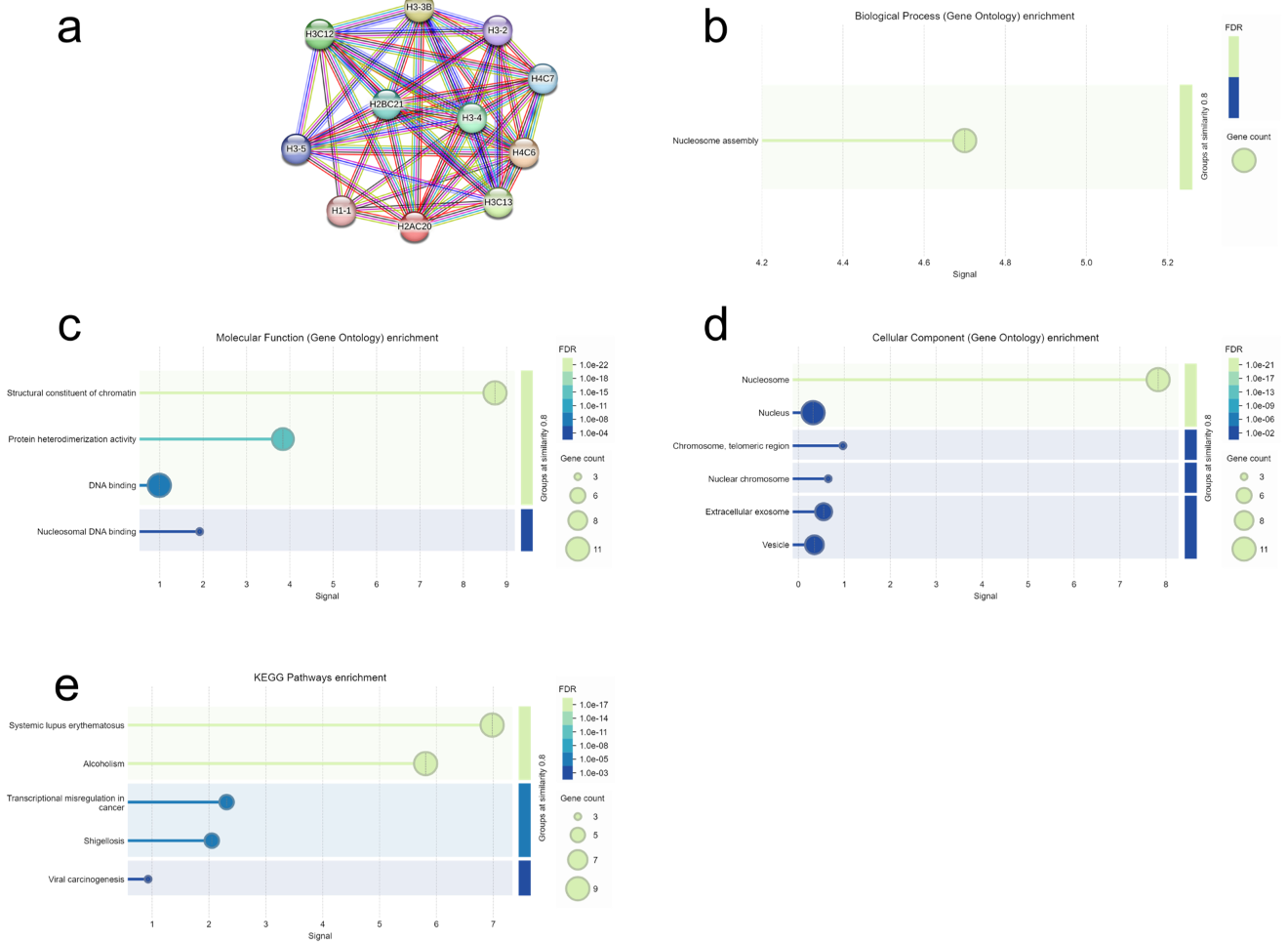

**Supplementary Figure 15. An interaction map for H3-3A and its functional cellular partners.** (a) protein-protein interaction map (b) Biological process (GO) enrichment (c) Molecular function (GO) enrichment (d) Cellular function (GO) enrichment (e) KEGG pathways enrichment. Protein interaction networks were generated using STRING database integrated with Cytoscape software (v3.10.2) with a minimum interaction score threshold of 0.4 for significantly differentially expressed proteins. Edges indicate protein-protein interactions, represented by different colors: known Interactions (Curated Databases: Light blue line, Experimentally Determined: Pink line), Predicted Interactions (Gene Neighborhood: Green line, Gene Fusions: Red line), Gene Co-occurrence: Blue line, Others (Text mining: Yellow line, Co-expression: Black line, Protein Homology: Lavender line).

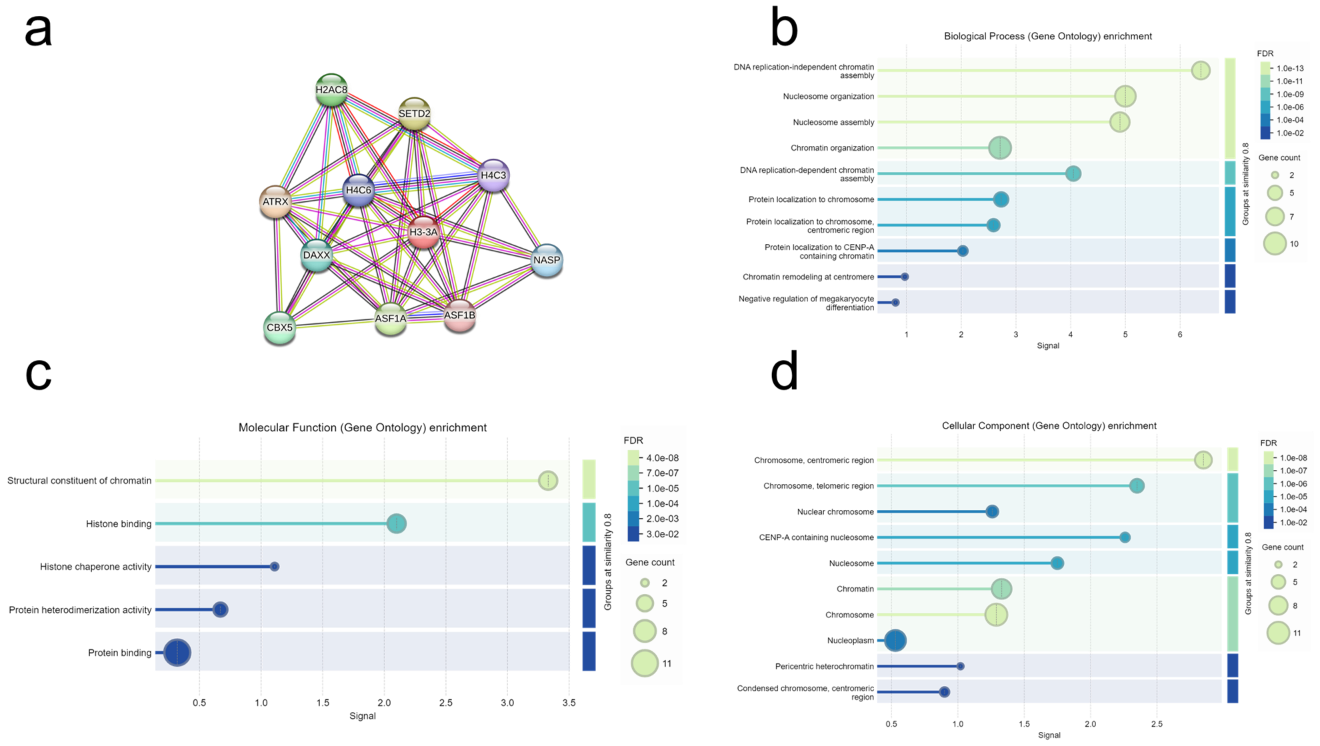

**Supplementary Figure 16. An interaction map for MGST1 and its functional cellular partners.** (a) protein-protein interaction map (b) Biological process (GO) enrichment (c) Molecular function (GO) enrichment (d) Cellular function (GO) enrichment (e) KEGG pathways enrichment. Protein interaction networks were generated using STRING database integrated with Cytoscape software (v3.10.2) with a minimum interaction score threshold of 0.4 for significantly differentially expressed proteins. Edges indicate protein-protein interactions, represented by different colors: known Interactions (Curated Databases: Light blue line, Experimentally Determined: Pink line), Predicted Interactions (Gene Neighborhood: Green line, Gene Fusions: Red line), Gene Co-occurrence: Blue line, Others (Text mining: Yellow line, Co-expression: Black line, Protein Homology: Lavender line).

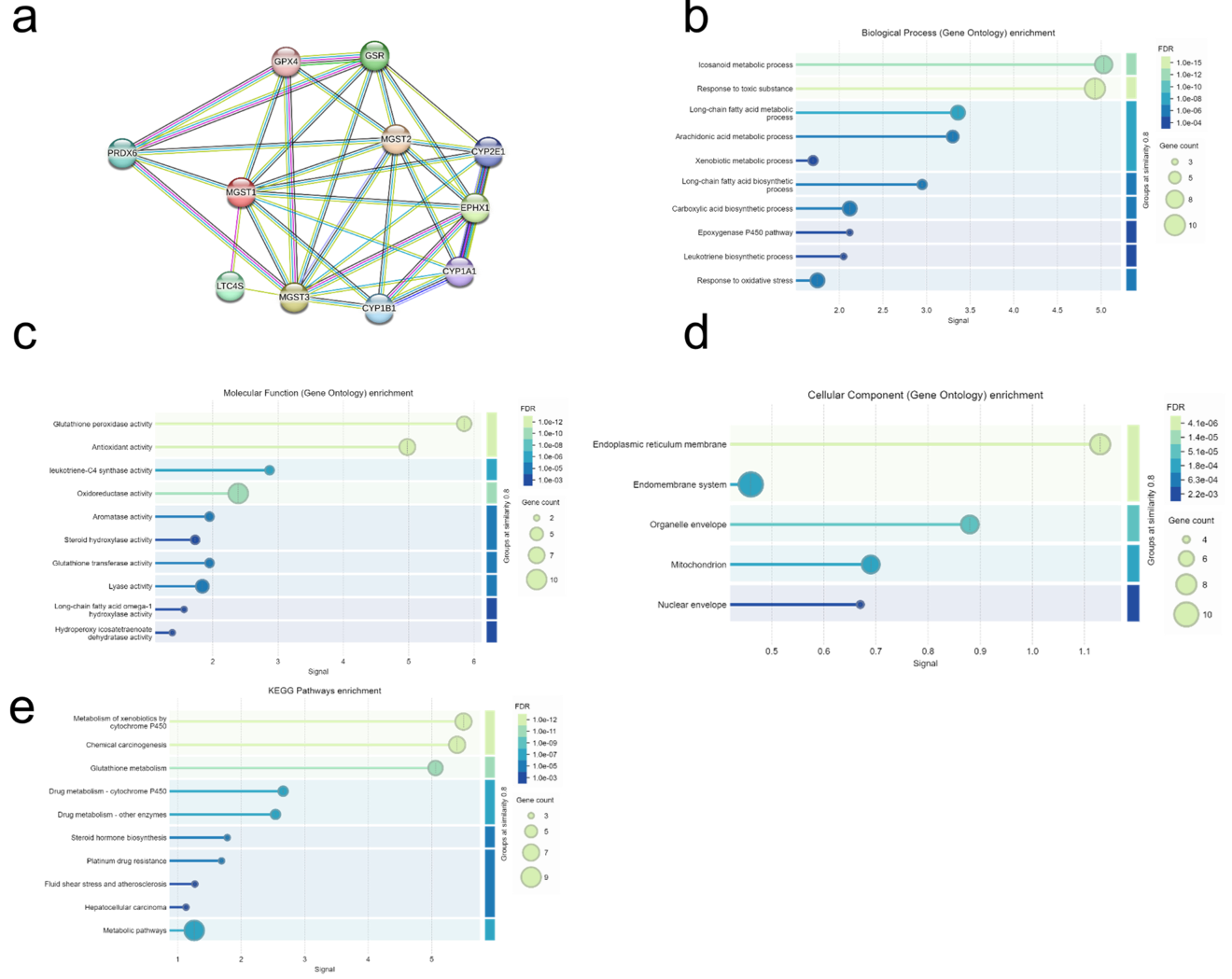

**Supplementary Figure 17. An interaction map for PSMB5 and its functional cellular partners.** (a) protein-protein interaction map (b) Biological process (GO) enrichment (c) Molecular function (GO) enrichment (d) Cellular function (GO) enrichment (e) KEGG pathways enrichment. Protein interaction networks were generated using STRING database integrated with Cytoscape software (v3.10.2) with a minimum interaction score threshold of 0.4 for significantly differentially expressed proteins. Edges indicate protein-protein interactions, represented by different colors: known Interactions (Curated Databases: Light blue line, Experimentally Determined: Pink line), Predicted Interactions (Gene Neighborhood: Green line, Gene Fusions: Red line), Gene Co-occurrence: Blue line, Others (Text mining: Yellow line, Co-expression: Black line, Protein Homology: Lavender line).

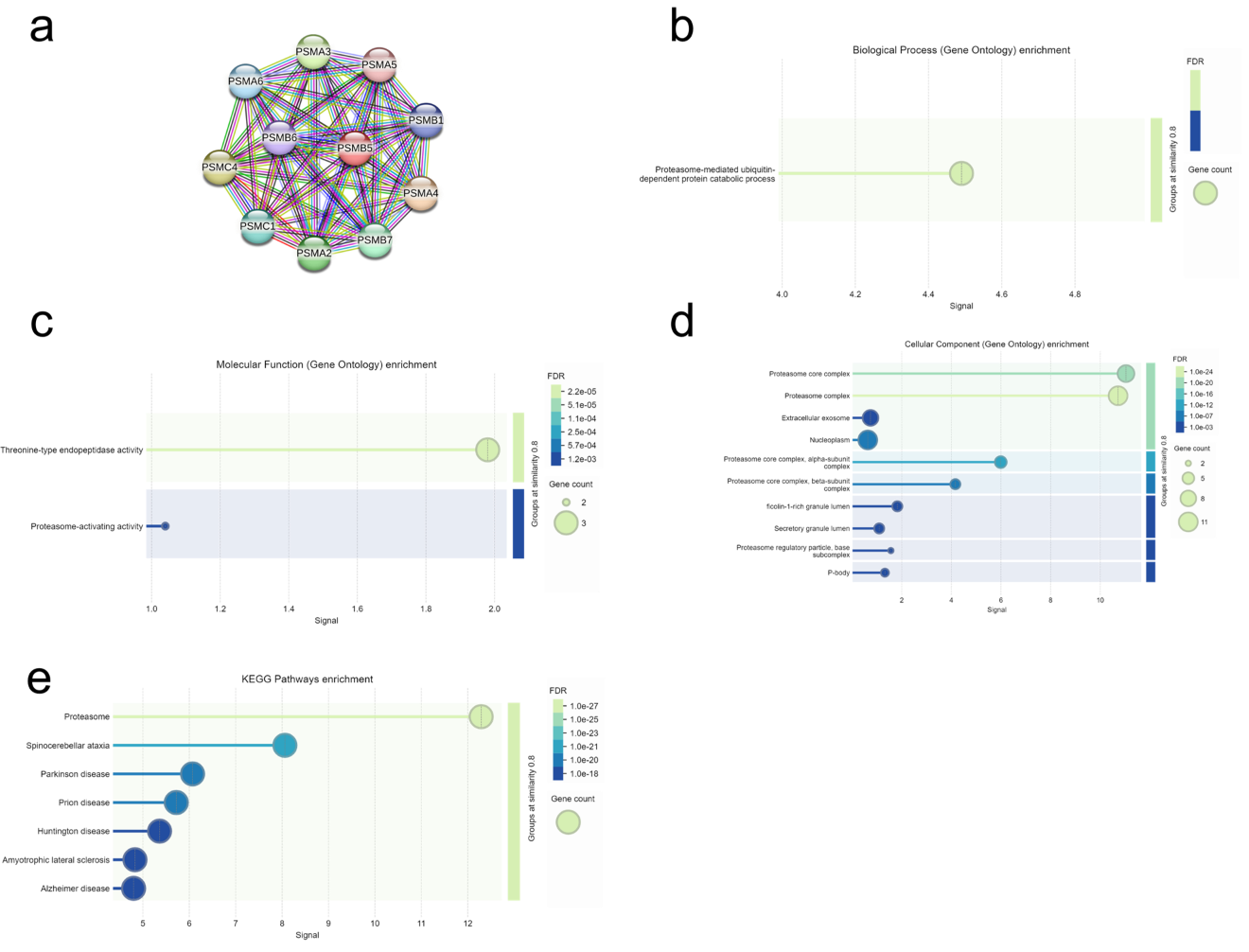

**Supplementary Figure 18. An interaction map for PSMA6 and its functional cellular partners.** (a) protein-protein interaction map (b) Biological process (GO) enrichment (c) Molecular function (GO) enrichment (d) Cellular function (GO) enrichment (e) KEGG pathways enrichment. Protein interaction networks were generated using STRING database integrated with Cytoscape software (v3.10.2) with a minimum interaction score threshold of 0.4 for significantly differentially expressed proteins. Edges indicate protein-protein interactions, represented by different colors: known Interactions (Curated Databases: Light blue line, Experimentally Determined: Pink line), Predicted Interactions (Gene Neighborhood: Green line, Gene Fusions: Red line), Gene Co-occurrence: Blue line, Others (Text mining: Yellow line, Co-expression: Black line, Protein Homology: Lavender line).

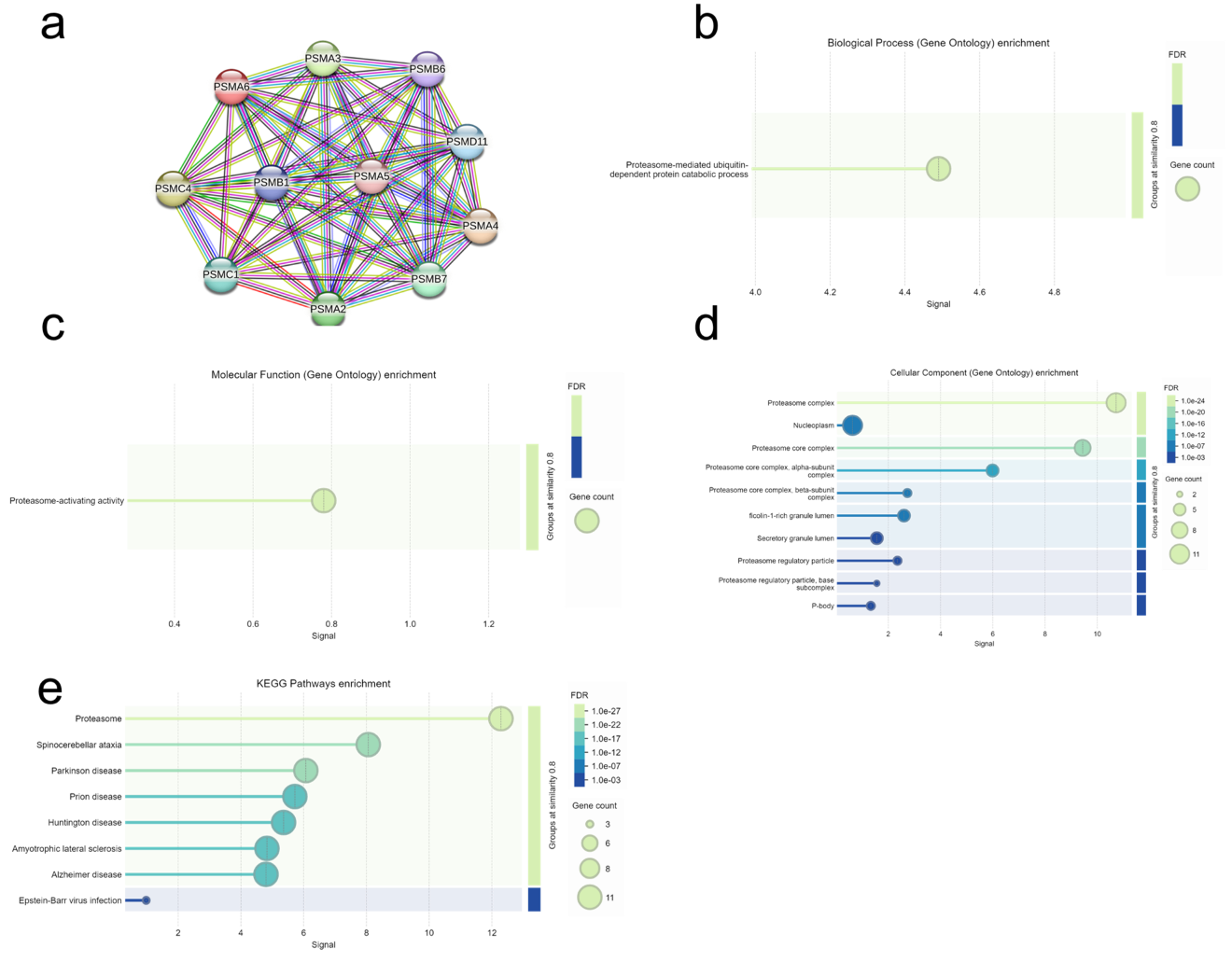

**Supplementary Figure 19. An interaction map for H4C6 and its functional cellular partners.** (a) protein-protein interaction map (b) Biological process (GO) enrichment (c) Molecular function (GO) enrichment (d) Cellular function (GO) enrichment (e) KEGG pathways enrichment. Protein interaction networks were generated using STRING database integrated with Cytoscape software (v3.10.2) with a minimum interaction score threshold of 0.4 for significantly differentially expressed proteins. Edges indicate protein-protein interactions, represented by different colors: known Interactions (Curated Databases: Light blue line, Experimentally Determined: Pink line), Predicted Interactions (Gene Neighborhood: Green line, Gene Fusions: Red line), Gene Co-occurrence: Blue line, Others (Text mining: Yellow line, Co-expression: Black line, Protein Homology: Lavender line).

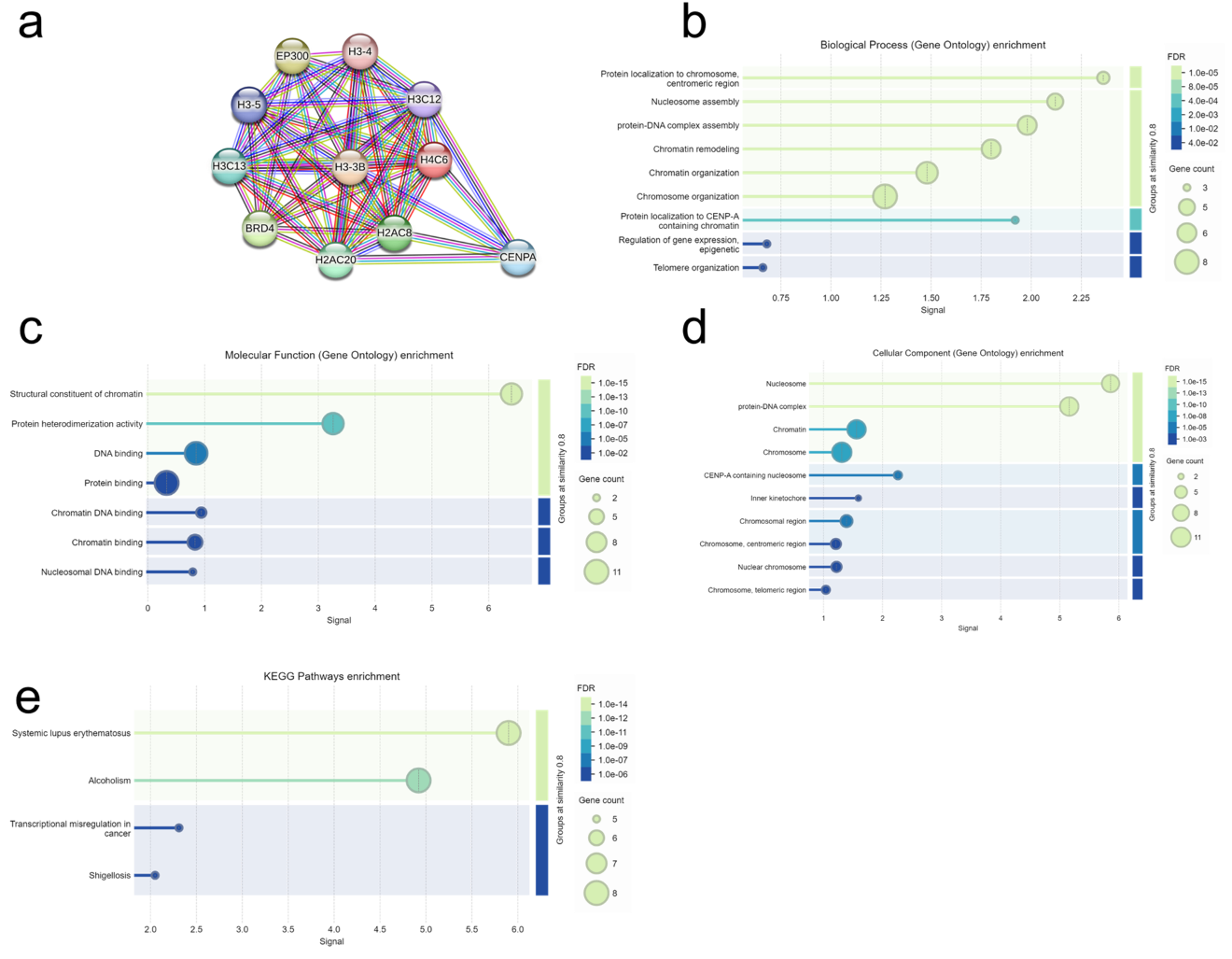

**Supplementary Figure 20. An interaction map for APOB and its functional cellular partners.** (a) protein-protein interaction map (b) Biological process (GO) enrichment (c) Molecular function (GO) enrichment (d) Cellular function (GO) enrichment (e) KEGG pathways enrichment. Protein interaction networks were generated using STRING database integrated with Cytoscape software (v3.10.2) with a minimum interaction score threshold of 0.4 for significantly differentially expressed proteins. Edges indicate protein-protein interactions, represented by different colors: known Interactions (Curated Databases: Light blue line, Experimentally Determined: Pink line), Predicted Interactions (Gene Neighborhood: Green line, Gene Fusions: Red line), Gene Co-occurrence: Blue line, Others (Text mining: Yellow line, Co-expression: Black line, Protein Homology: Lavender line).

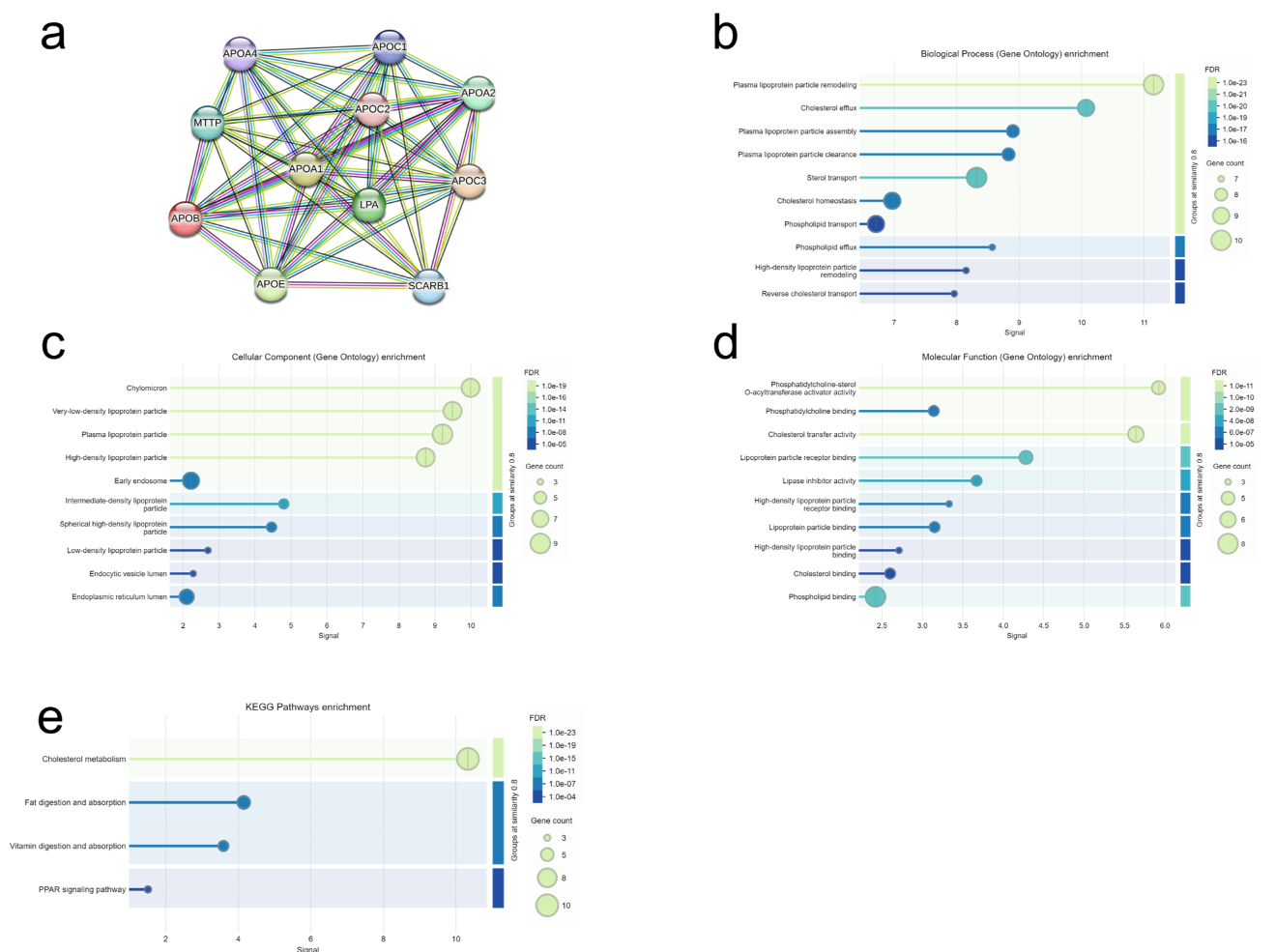

**Supplementary Figure 21. An interaction map for APCS and its functional cellular partners.** (a) protein-protein interaction map (b) Biological process (GO) enrichment (c) Molecular function (GO) enrichment (d) Cellular function (GO) enrichment (e) KEGG pathways enrichment. Protein interaction networks were generated using STRING database integrated with Cytoscape software (v3.10.2) with a minimum interaction score threshold of 0.4 for significantly differentially expressed proteins. Edges indicate protein-protein interactions, represented by different colors: known Interactions (Curated Databases: Light blue line, Experimentally Determined: Pink line), Predicted Interactions (Gene Neighborhood: Green line, Gene Fusions: Red line), Gene Co-occurrence: Blue line, Others (Text mining: Yellow line, Co-expression: Black line, Protein Homology: Lavender line).

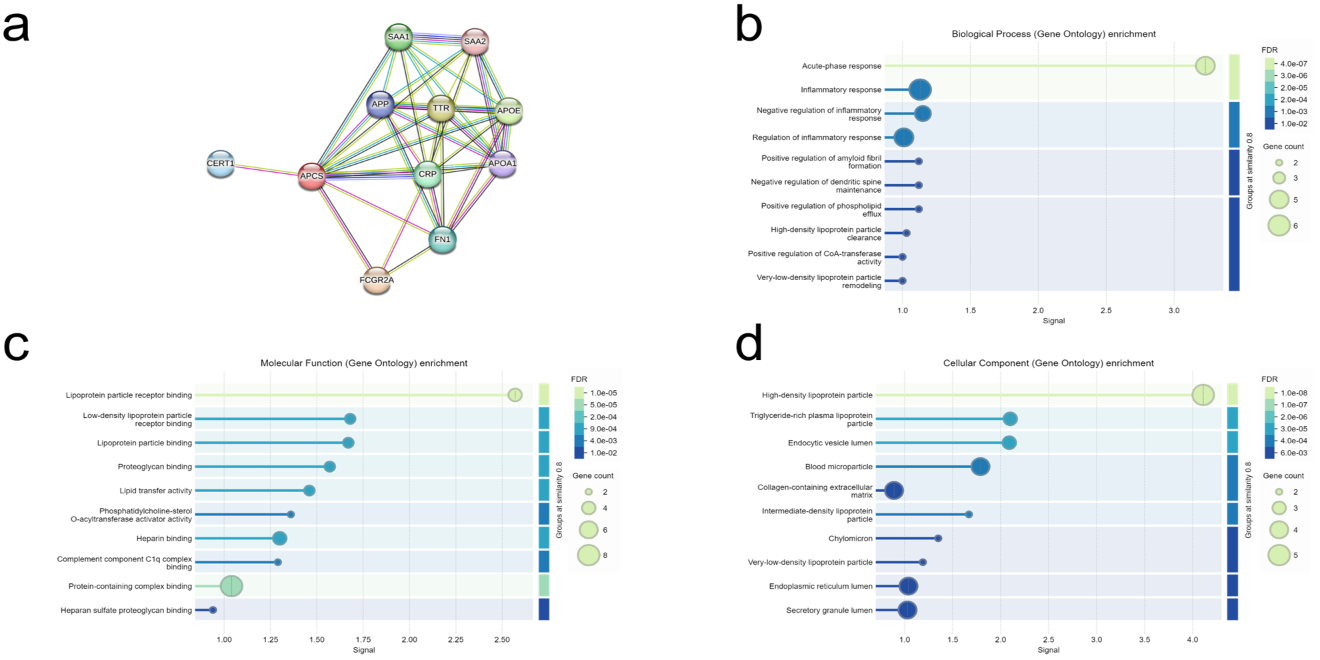

**Supplementary Figure 22. An interaction map for CD36 and its functional cellular partners.** (a) protein-protein interaction map (b) Biological process (GO) enrichment (c) Molecular function (GO) enrichment (d) Cellular function (GO) enrichment (e) KEGG pathways enrichment. Protein interaction networks were generated using STRING database integrated with Cytoscape software (v3.10.2) with a minimum interaction score threshold of 0.4 for significantly differentially expressed proteins. Edges indicate protein-protein interactions, represented by different colors: known Interactions (Curated Databases: Light blue line, Experimentally Determined: Pink line), Predicted Interactions (Gene Neighborhood: Green line, Gene Fusions: Red line), Gene Co-occurrence: Blue line, Others (Text mining: Yellow line, Co-expression: Black line, Protein Homology: Lavender line).

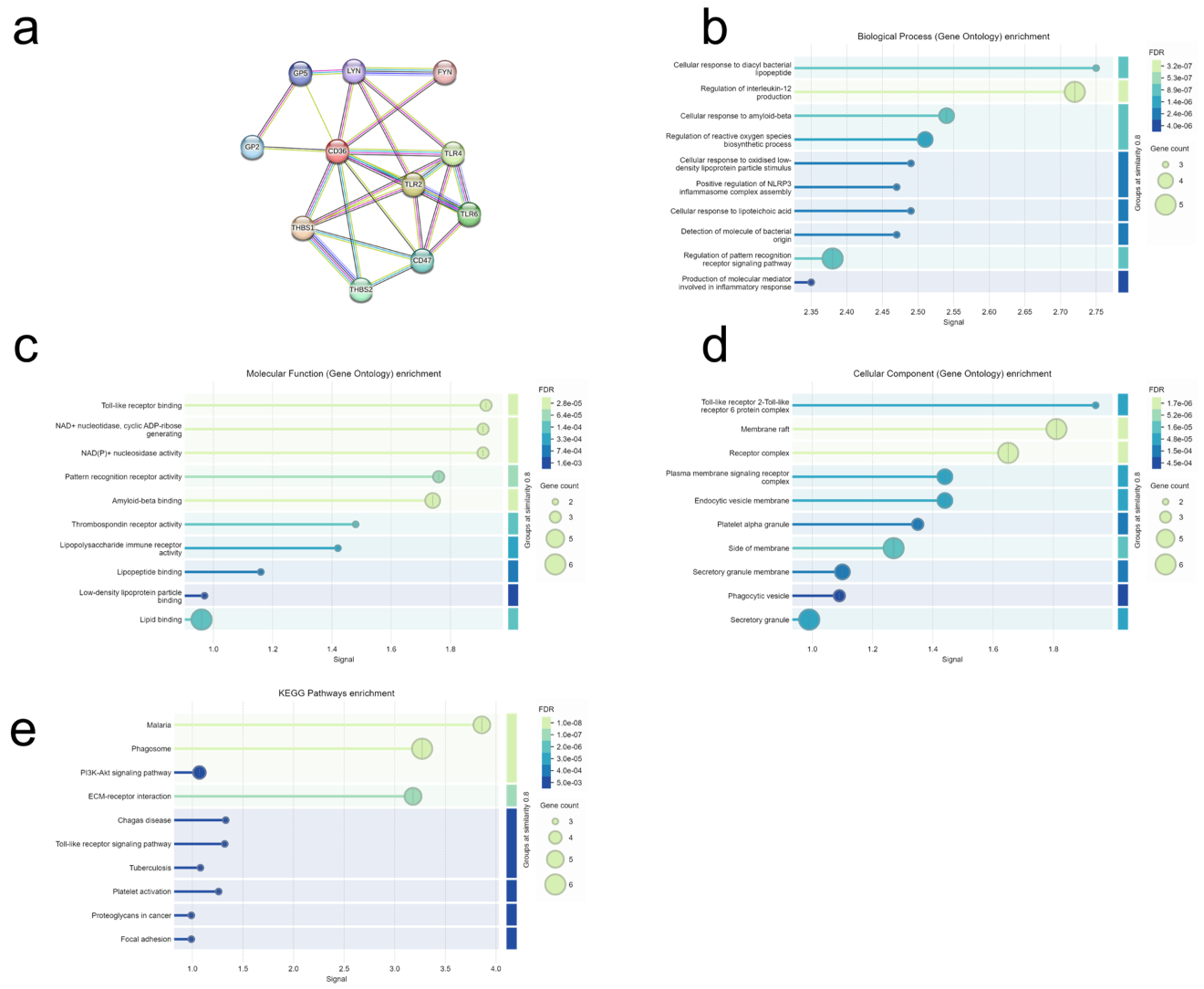

**Supplementary Figure 23. An interaction map for CES1 and its functional cellular partners.** (a) protein-protein interaction map (b) Biological process (GO) enrichment (c) Molecular function (GO) enrichment (d) Cellular function (GO) enrichment (e) KEGG pathways enrichment. Protein interaction networks were generated using STRING database integrated with Cytoscape software (v3.10.2) with a minimum interaction score threshold of 0.4 for significantly differentially expressed proteins. Edges indicate protein-protein interactions, represented by different colors: known Interactions (Curated Databases: Light blue line, Experimentally Determined: Pink line), Predicted Interactions (Gene Neighborhood: Green line, Gene Fusions: Red line), Gene Co-occurrence: Blue line, Others (Text mining: Yellow line, Co-expression: Black line, Protein Homology: Lavender line).

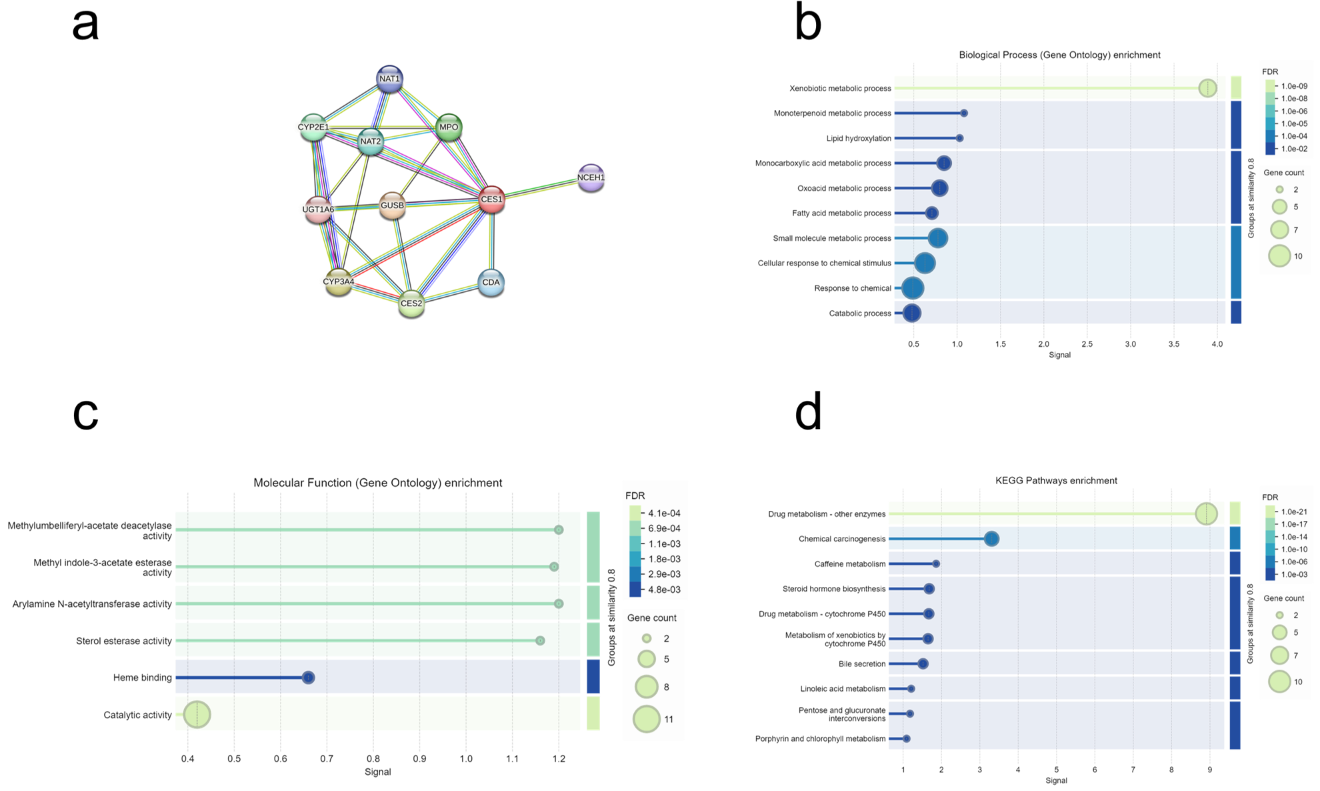
